# Decision Tree Ensembles Utilizing Multivariate Splits Are Effective at Investigating Beta-Diversity in Medically Relevant 16S Amplicon Sequencing Data

**DOI:** 10.1101/2022.03.31.486647

**Authors:** Josip Rudar, G. Brian Golding, Stefan C. Kremer, Mehrdad Hajibabaei

## Abstract

Developing an understanding of how microbial communities vary across conditions is an important analytical step. We used 16S rRNA data isolated from human stool to investigate if learned dissimilarities, such as those produced using unsupervised decision tree ensembles, can be used to improve the analysis of the composition of bacterial communities in patients suffering from Crohn’s Disease and adenomas/colorectal cancers. We also introduce a workflow capable of learning dissimilarities, projecting them into a lower dimensional space, and identifying features that impact the location of samples in the projections. For example, when used with the centered log-ratio transformation, our new workflow (TreeOrdination) could identify differences in the microbial communities of Crohn’s Disease patients and healthy controls. Further investigation of our models elucidated the global impact ASVs had on the location of samples in the projected space and how each ASV impacted individual samples in this space. Furthermore, this approach can be used to integrate patient data easily into the model and results in models that generalize well to unseen data. Models employing multivariate splits can improve the analysis of complex high-throughput sequencing datasets since they are better able to learn about the underlying structure of the dataset.

**Author Summary:** There is an ever-increasing level of interest in accurately modeling and understanding the role that commensal organisms play in human health and disease. We show that learned representations can be used to create informative ordinations. We also demonstrate that the application of modern model introspection algorithms can be used to investigate and quantify the impact of taxa in these ordinations and that the taxa identified by these approaches have been associated with immune-mediated inflammatory diseases and colorectal cancer.

## Introduction

The analysis of the composition of ecological communities is essential to determine their function within the broader environment and within host. The statistical analysis of differences between communities, beta diversity, is often an important aspect of statistical pipelines used to investigate ecological and metagenomic datasets. Central to the analysis of beta diversity is the calculation of pairwise distances or dissimilarities between samples. This calculation is carried out by carefully choosing and then applying a particular dissimilarity which is known to measure an important characteristic of the data. For example, some methods (such as correlation-based distance methods) perform well at measuring environmental gradients while others, such as the Jaccard distance, can perform well when one is interested in clustering data (1–3). When analyzing amplicon sequencing data another typical goal is to discover amplicon sequence variants (ASVs) or operational taxonomic units (OTUs) associated with each type of community. This task is inextricably linked to variations in the composition of communities, which can be captured using a distance metric or dissimilarity measure. This type of analysis is typically carried separately from the analysis of community composition using tools such as DESeq2, MetagenomeSeq, ANCOM, or LEfSE (3–8). Furthermore, the assumptions of these methods could contribute to higher false-positive and false-negative rates which could lead to potentially erroneous or incomplete interpretations (3,9,10). Therefore, it is preferable if impactful ASVs (or OTUs, etc) can be identified directly when investigating community composition. Furthermore, it is also important that these analyses are done using methods that make few (if any) assumptions about the underlying distribution of each ASV. Finally, for any method to be useful, it is important for it to consider dependencies between features (which can be genes, taxonomic groups, operational taxonomic units, amplicon sequence variants). By attempting to model these dependencies, subtle differences between groups are more likely to be detected. This is supported by recent work which has demonstrated that genomic, transcriptomic, and metagenomic datasets are better understood if these relationships are considered (11–13).

Machine learning algorithms are uniquely suited to address these challenges. Unlike statistical models, machine learning models tend not to assume anything about the underlying distribution of each feature (4,5). Furthermore, some machine learning models, such as Random Forests (RFs) and related classifiers, are capable of identifying dependencies between features without the need for the user to explicitly include these dependencies in the model (11,14–17). One, and arguably underused, ability inherent to this class of models is that they can be used in an ‘unsupervised’ manner to learn a dissimilarity function (15,18,19). This is known as metric learning and the learned dissimilarity function can be used to replace a more traditional method (such as the Jaccard distance or Bray-Curtis dissimilarity) when investigating beta-diversity (17,20). This approach is also advantageous since it learns to remove the influence of uninformative features (21). Unsupervised Random Forests have previously been used to discover similar cell populations in single-cell RNAseq data, identify different classes of renal cell carcinomas tumors, and study the underlying structure of a population using shared genetic variations (20,22,23). If this approach is applied to amplicon sequencing data it may be possible to simultaneously visualize the differences between communities while also identifying which features contribute most to the placement of each community within the projected space (17,20). However, Random Forests do suffer an important limitation: the decision trees used to construct the forest make axis-orthogonal cuts. Randomizing the selection of cut points at each node has been shown to help since this results in the construction of better decision boundaries. However, improvements such as these are not a solution since axis-orthogonal cuts are still made. To solve this problem trees that can learn oblique or non-linear cuts should be used. Recent and historical work using decision tree classifiers demonstrates that these types of splits often result in learning a more appropriate representation of the data (11,24).

Recently, we have introduced a new algorithm, Large Scale Non-Parametric Discovery of Markers (LANDMark), as an alternative to RFs (25). Like the RF, LANDMark is a tree-based approach that recursively partitions samples until a stopping criterion is met. Unlike RFs, LANDMark can use learned linear and non-linear models to partition samples (25). In this study, we investigate LANDMark’s ability to predict medically relevant outcomes, such as Crohn’s Disease and colorectal cancer, using amplicon sequence datasets. We also provide a workflow, TreeOrdination, which uses LANDMark to quantify differences in the makeup of microbial communities in an unsupervised manner. TreeOrdination and LANDMark’s performance is investigated using a synthetic and randomized dataset, a small case study, and a larger colorectal cancer dataset (26–28). The code used to perform the analysis in this work can be found at: https://github.com/jrudar/Unsupervised-Decision-Trees.

## Methods

### Dataset Description

Two human microbiome datasets were selected for use as case studies. The first dataset was chosen since it contains samples from patients who suffer from immune-mediated inflammatory diseases (IMID) (26). Differences between the microbiomes of patients suffering from Crohn’s disease (CD), ulcerative colitis (UC), multiple sclerosis (MS), and rheumatoid arthritis (RA) were compared to healthy controls. This is an important area of research since many people across the world suffer from these conditions. Therefore, models which are better able to identify ASVs associated with these conditions can provide further insight into these diseases. In this study, we used the CD-HC subset of data as a case study. The second dataset is derived from the stool samples of healthy patients and those suffering from colorectal cancers and adenomas. 16S rRNA gene sequencing was used to identify the composition of the colon community in each of the 490 patients (28). This dataset is important due to the prevalence of colorectal cancers in our society. Therefore, the ability to train models to detect the early stages of cancer development and colorectal cancers will result in economic savings and the saving of lives.

### Bioinformatic Processing of Raw Reads

Raw sequences for each of the datasets were obtained from the Sequence Read Archive (PRJNA450340 and SRP062005) (26,28). All bioinformatic processing of the raw reads was prepared using the MetaWorks v1.8.0 pipeline (available online at: https://githib.com/terrimporter/MetaWorks) (29). The default settings for merging reads were used except for the parameter controlling the minimum fraction of matching bases, which was increased from 0.90 to 0.95. This was done to remove a larger fraction of potentially erroneous reads. Merged reads were then trimmed using the default settings MetaWorks passes to CutAdapt. Since reads from SRP062005 were pre-processed and the primers removed no reads were discarded during trimming. The remaining quality-controlled sequences were then de-replicated and denoised using VSEARCH version 2.15.2 to remove putative chimeric sequences (30). Finally, VSEARCH was used to construct a matrix where each row is a sample and each column an Amplicon Sequence Variant (ASV). Taxonomic assignment was conducted using the RDP Classifier (version 2.13) and the built-in reference set (31). ASVs that are likely to be contaminants, specifically those likely belonging to chloroplasts and mitochondria, were removed. From the remaining sequences, only those belonging to the domain *Bacteria* and *Archaea* were retained for further analysis. A sample by ASV count matrix, where each row is a sample and each column an ASV, was created for each dataset.

### Calculation of Dissimilarities

Pairwise dissimilarities between samples were calculated using the Jaccard distance for presence-absence data (Equation 1), the Aitchison distance for centered-log and robust centered-log ratio data (Equation 2), and the Bray-Curtis dissimilarity (Equation 3) for data converted into proportions. In addition, we investigated the performance of Extremely Randomized Trees dissimilarity (Equations 4 and 5), and LANDMark dissimilarity. In these equations *X*_*i*_ and *X*_*j*_ represent the *i*^*t*h^ and *j*^th^ samples in *X*. Distances were calculated using the ‘pairwise_distances’ function from scikit-learn (version 1.1.2) (32).

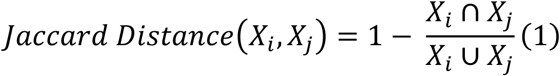

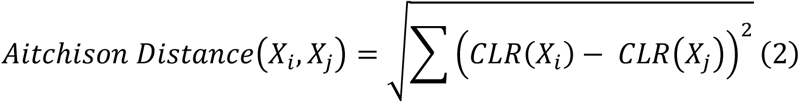

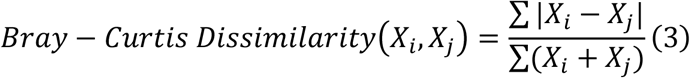

To calculate the Extra Trees dissimilarity, one simply determines the label of each terminal leaf into which a sample falls. This is recorded for each decision tree in the forest. These labels are then used to construct a binary matrix, 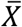, which simply describes the terminal leaves into which each sample falls. Each row of 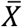 is a sample and each column is a leaf label. The similarity between two samples is then found by calculating how often two samples share a terminal node and dividing this count by the number of trees in the forest, *N* (Equation 4). Finally, the similarity is turned into a dissimilarity according to Equation 5 (19,20,22,33).

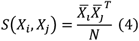

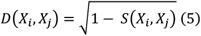

LANDMark returns this binary representation directly (25). To calculate the dissimilarity between samples in the LANDMark embedding, the Hamming distance can be used since we are only interested in the positions where leaf labels are not shared. In these experiments, we used LANDMark version 1.2.2 and the implementation of the Extra Trees classifier found in scikit-learn (25,32).

To use decision tree ensembles in an unsupervised manner a second randomized dataset is needed. To create the randomized dataset, a copy of the original count matrix was created. Then, the counts of each ASV in the copy are randomly sampled without replacement (19,34). The original data and randomized copy are then combined. To prevent Extra Trees and LANDMark from using information about the original class labels, the rows in the randomized dataset are assigned a label of “Random” while original samples are assigned a label of “Original”. The combined dataset and the new labels are then used to train the decision-tree ensemble. Dissimilarities are calculated by passing the original data to the model to determine which terminal leaves are associated with each sample.

### TreeOrdination: Automating the Identification of Informative ASVs When Using Tree Dissimilarities

We have developed a workflow, TreeOrdination, which uses a relatively simple set of steps to create and analyze projections of high-dimensional amplicon sequencing data. This was done to investigate if tree-based dissimilarities can be used to investigate beta diversity. The TreeOrdination and LANDMark source code and documentation are available at https://github.com/jrudar/LANDMark and https://github.com/jrudar/TreeOrdination.

TreeOrdination is a wrapper around the LANDMark and Uniform Manifold Approximation and Projection (UMAP) algorithms (25,35). The default settings for LANDMark were used except for the number of trees (which was set to 160), the ‘use_nnet’ parameter (which was set to ‘False’), and the ‘max_samples_tree’ parameter (which was set to 100) (25). Upon completion of training, the original training data is passed through the classifier and the binary matrix describing which terminal leaves each sample falls into is extracted. Multiple LANDMark representations can be created and concatenated together. These representations can be used to train an Extra Trees classifier to predict the original class labels. In addition, the LANDMark representation is projected into a smaller 15-dimensional space using Uniform Manifold Approximation and Projection (UMAP) version 0.5.3 (35,36). To create this projection the ‘min_dist’ parameter is set to 0.001, while the ‘n_neighbors’ and ‘metric’ parameters are set to 8 and “hamming”, respectively. The ‘PCA’ functionality of scikit-learn is then used to summarize the major directions of variation in UMAP representation (Supplementary Figure 1) (32). To understand which ASVs impact the location of samples within the PCA projection, the original data is then used to train an Extra Trees regression model to predict the location of each sample in the projected space. This is necessary since calculating the impact that each ASV has on the location of each sample in the projected space using the original model is a computationally expensive task. Shapley scores are then used to calculate global feature importance scores and per-sample importance scores, respectively (37,38). This method of investigating black-box models uses game theory to calculate the importance and contribution of each feature to the final prediction. This is done calculating how often a feature contributes to a prediction when included with a coalition of other features.

### Creation of Positive and Negative Controls

Positive and negative controls were created using a copy of the code found at https://github.com/cameronmartino/deicode-benchmarking/blob/master/simulations (27). This code was used in the simulations which tested robust centered log-ratio transformation introduced by Martino et al. (2019) (27). Briefly, a synthetic dataset containing 200 samples and 1000 features was constructed according to the procedure found by Martino et al. (2019). This was a positive control. A copy of this dataset was made, and each feature column was independently permuted to create the negative control.

### Analysis of Beta-Diversity and Generalization Performance in the Case Studies and Synthetic Data

Samples were split into training and testing sets using repeated stratified 5-fold cross-validation. Five repeats were used. ASVs occurring in two or fewer training samples (in the case studies) and 5% of training samples (in the case of the control datasets) were identified. These ASVs were then removed from both the training and testing samples. This filtration step was taken since a reduction in the number of features can often lead to a more generalizable model (39–41). Following this, the size of each training library was calculated, and the 15^th^ percentile of these sizes was found. Training and testing libraries were then rarefied so that each library was this size and those smaller than the 15^th^ percentile were removed (7). The rarefied data was used for tests involving the presence-absence and proportion transformations. Tests using rCLR and CLR transformed data did not involve rarefaction as these transformations can naturally handle compositions (6,27,42,43). Count matrices were CLR and rCLR transformed using the scikit-bio (version 0.5.7) and DEICODE (version 0.2.4) packages, respectively (27,44,45). To produce the rCLR transformed data 1000 iterations of DEICODE’s matrix completion algorithm was used. Following this, the *U, V*, and *S* matrices (which are analogous to the sample loading, feature loading, and eigenvalue matrix of singular value decomposition) were extracted for further analysis (27). Randomized data for training unsupervised tree models were also created as described above and the presence-absence, proportion, CLR, and rCLR transformations were applied.

The transformed training data was used as input for a PerMANOVA analysis (using 999 replications). The training data was also projected using principal coordinates analysis (PCoA) /Robust Principal Components Analysis (RPCA) and UMAP to create two-dimensional projections which were used as input into another PerMANOVA analysis (using 999 replications). In this experiment, high F-statistics associated with significant p-values (defined as p ≤ 0.05) provide evidence that differences exist between groups and larger F-statistics provide evidence for a better separation between groups. PCoA projections were created scikit-bio while UMAP projections were created using the ‘umap’ package (35,44). In the case of rCLR transformed data, this projection was automatically supplied in the form of the *U* matrix. Unsupervised Extremely Randomized Trees and Unsupervised LANDMark projections were created by training an Extra Trees and LANDMark Classifier. Both ensembles were consutructed using 160 estimators. The ‘use_nnet’ and ‘max_samples_tree’ parameters of the LANDMark classifier were set to ‘False’ and ‘100’. Dissimilarities calculated using the unsupervised ensembles and the PCoA and UMAP projections of these dissimilarities were used as input into a PerMANOVA analysis. Finally, TreeOrdination ordinations (using 160 estimators, 5 LANDMark classifiers, and 100 samples per tree) were created. These ordinations were also used as input into a PerMANOVA analysis.

Where appropriate, the transformed and projected training data was also used to train an Extra Trees and LANDMark classifier. The high-dimensional TreeOrdination embedding was used to train an ExtraTrees classifier. The generalization performance of each model was measured using the balanced accuracy score (from scikit-learn) (32). This was chosen since it accounts for differences in class sizes when working with unbalanced datasets. To ascertain which models performed the best a Wilcoxon Signed Rank test followed by the Benjamini-Hochberg correction, implemented in the Python ‘statannotations’ package (version 0.5), was used (46).

To investigate which ASVs are different between sample types, we construct a TreeOrdination model (as described above) using the Crohn’s Disease data. For this model, 70% of the samples are used for training data while the remaining 30% are used for testing. Shapley scores for each group were calculated to determine which features impact the location of samples in ordination space (28,37,38). The projection was visualized using the Python ‘seaborn’ (version 0.11.2) and ‘matplotlib’ (version 3.5.2) packages (47,48)

### Analysis of Generalization Performance in the Colorectal Cancer Data

It is important to investigate how TreeOrdination and LANDMark behave when faced with data where the differences between classes are not as clear. To carry out this investigation, we used a large colorectal cancer dataset consisting of 490 samples (28). Since the goal of this experiment is to determine how well models which make multivariate cuts perform when the effect size between groups is smaller, a simplified analysis was conducted. The data was split into two groups: normal vs colorectal cancer and normal vs. lesion (adenomas and colorectal cancers). The generalization performance using the ASV data and ASV data complemented using the fecal immunochemical test (FIT) was then measured (28). Five-fold stratified cross-validation with five repeats was used to split the samples from each group into training and testing data. ASVs found in fewer than 5% of samples in the training data were identified and these ASVs were removed from both training and testing data. The counts for each sample were transformed into frequencies, with zero counts being replaced using the multiplicative replacement procedure and the centered-log-ratio transformation was then applied (45). A random classifier (using a stratified sampling strategy), Extra Trees classifier (using 160 trees), Random Forest classifier (using 160 trees), LANDMark classifier (using 160 trees with each tree randomly selecting 100 samples, and the ‘use_nnet’ parameter set to False), and TreeOrdination (using 160 trees with each tree randomly selecting 100 samples) were used to measure generalization performance. The balanced accuracy and ROC-AUC scores were then calculated on each fold. The Wilcoxon Signed Rank Test followed by the Benjamini-Hochberg correction for multiple comparisons was used to determine if the generalization performance between models was equivalent (46).

For each test (colorectal cancer vs normal, lesion vs normal) we measured the ability of LANDMark and TreeOrdination to identify colorectal cancers and lesions. This was done by first determining the probability threshold which maximizes the balanced accuracy score of the model concerning the prediction of lesions. To do this we split each subset into training and testing folds using five-fold stratified cross-validation with five repeats as described above. After training each model, the probability of colorectal cancer was determined using the test set. These probabilities were then averaged across folds. A list of probability thresholds (spaced evening from 0 to 1.0 using increments of 0.01) was then used to determine the best possible cutoff for colorectal cancer vs normal and the normal vs lesion models. Finally, we investigated which ASVs were different between sample types by constructing a TreeOrdination model (as described above). For this model, 80% of the samples were used for training data while the remaining 20% were used for testing. Prediction probabilities were calculated using TreeOrdination. The subset of adenoma and colorectal cancer samples at or above this threshold were extracted and normal samples below this threshold in the test set were identified. These samples were then grouped into FIT positive and FIT negative samples and plotted. Shapley scores for each group were calculated to determine which features impact the location of samples in ordination space (28,37,38).

## Results

### TreeOrdination Projections Result in Well Separated Groups in Simulated Data

Under each transformation using the negative controls, we observed low F-statistics and non-significant p-values. This observation occurred irrespective of ordination type and model. PerMANOVA was able to identify significant differences (p ≤ 0.001) between groups found in the positive controls across each cross-valiation fold (Figure 1). UMAP, Unsupervised LANDMark, and TreeOrdination being among the best-performing methods (Figure 1). However, the sole use of F-statistics (or an equivalent statistic) to demonstrate class separation in the projection is not enough. Each ordination method should also be able to place new data of the same class into a similar location in the ordination space. When running this experiment, however, it quickly became apparent that creating ordinations using PCoA or RPCA results in an important limitation. Specifically, both methods are unable to transform and project new data into the space created using the training data. To do this, we would have needed to use all the available data (train, test) and this would have resulted in data leakage since the test data would influence the final projection. Therefore, to prevent data leakage and the reporting of overly optimistic and potentially misleading results, the generalization performance of RPCA and PCoA ordination methods were excluded (49). An overview of the important properties of each ordination method is presented in Table 1. Of the remaining transformations, both LANDMark, Extra Trees, and TreeOrdination models were successful at classifying unseen samples (Figure 2). Although statistically significant differences between models were observed, the size of these differences is likely small in this test since all classifiers resulted in nearly perfect classification results while balanced accuracy scores (Supplementary Figure 2). In contrast, when training models using the negative control data, we observed random performance (Figure 3). Finally, our results show that projecting data using UMAP resulted in good overall performance regardless of distance or dissimilarity measure.

**Figure 1:**
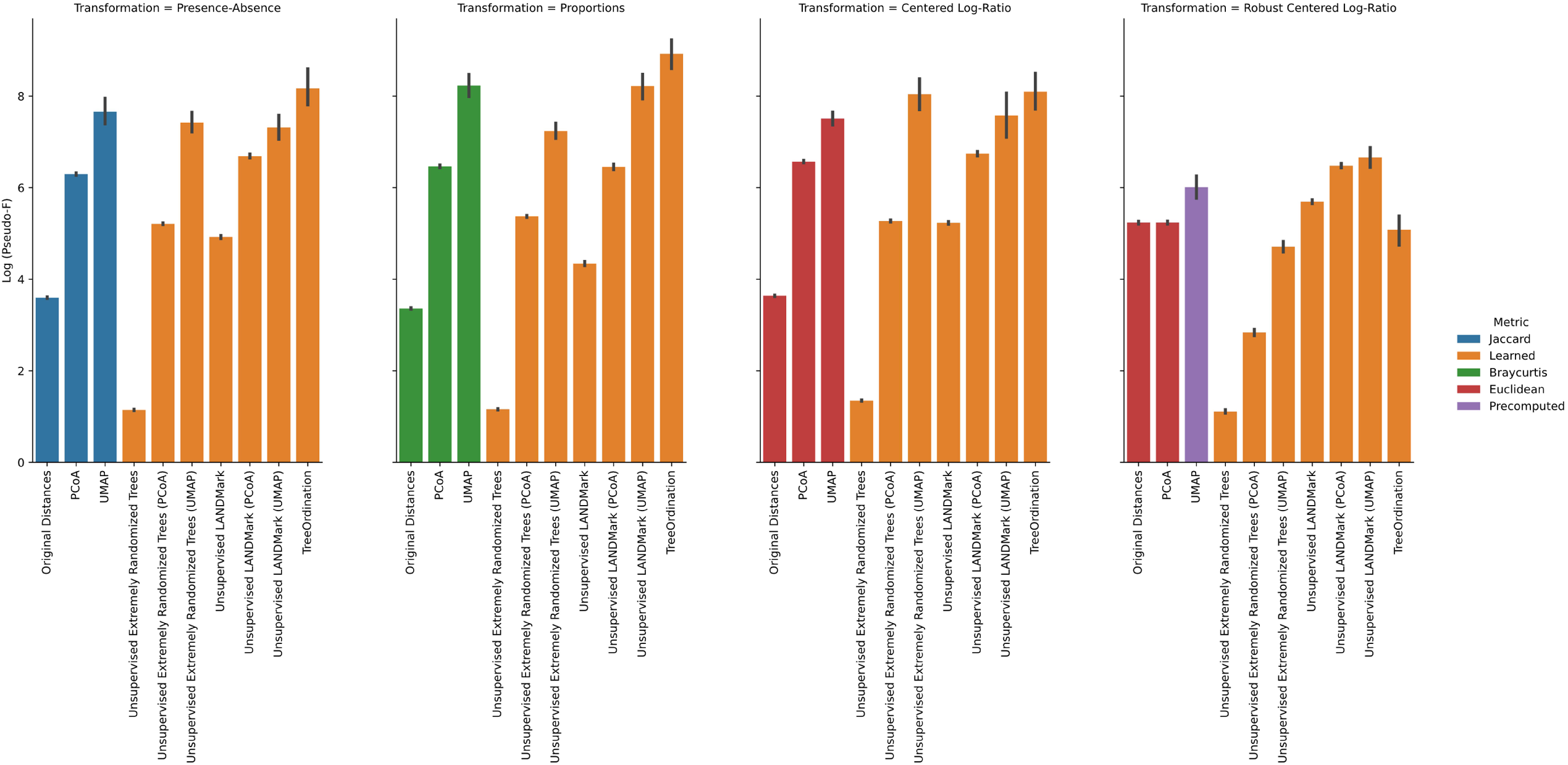
PerMANOVA results for each test condition in the positive control data. The pseudo-F statistic is log-transformed to improve the readability of the chart. Higher statistics indicate a greater separation between groups of samples. The 95% confidence interval, calculated using 2000 bootstraps, is shown for each bar in black. Five-fold stratified cross-validation with five repeats was used to generate this data. Bars are colored according to the dissimilarity. Blue, green, and red bars represent data analyzed using the Jaccard distance, Bray-Curtis dissimilarity, and Euclidean distances. Orange bars are learned dissimilarities, while the purple bar are distances calculated using DEICODE.

**Table 1:**
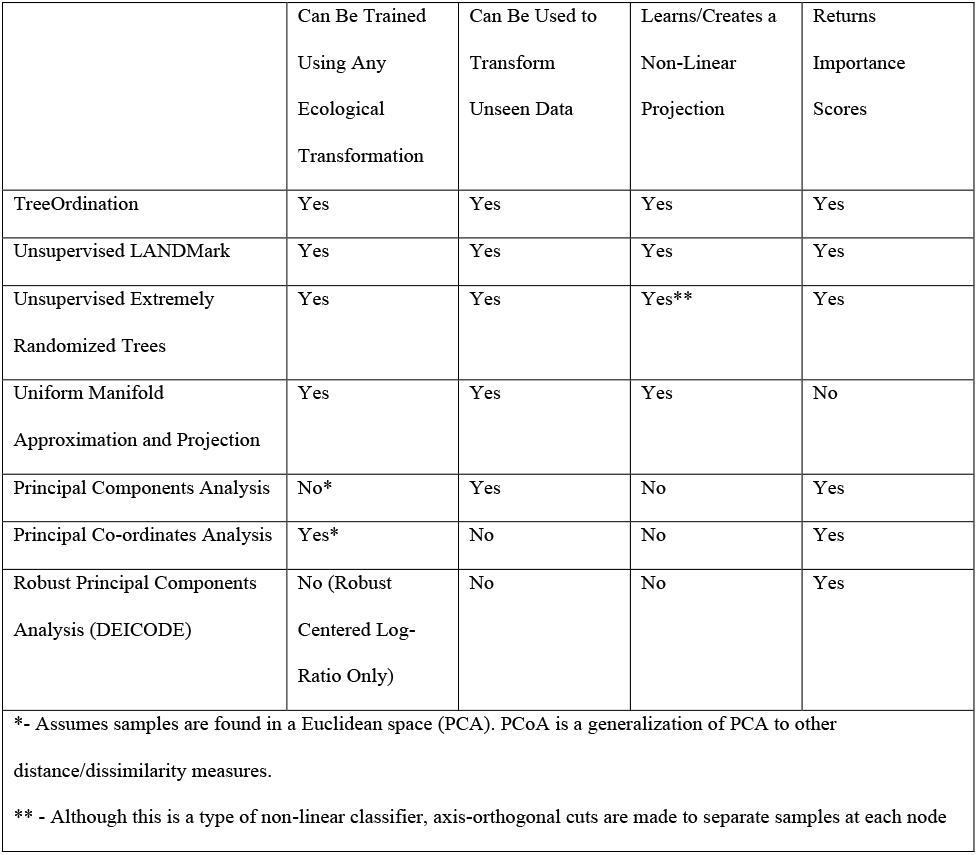
An overview of the properties of each ordination method.

**Figure 2:**
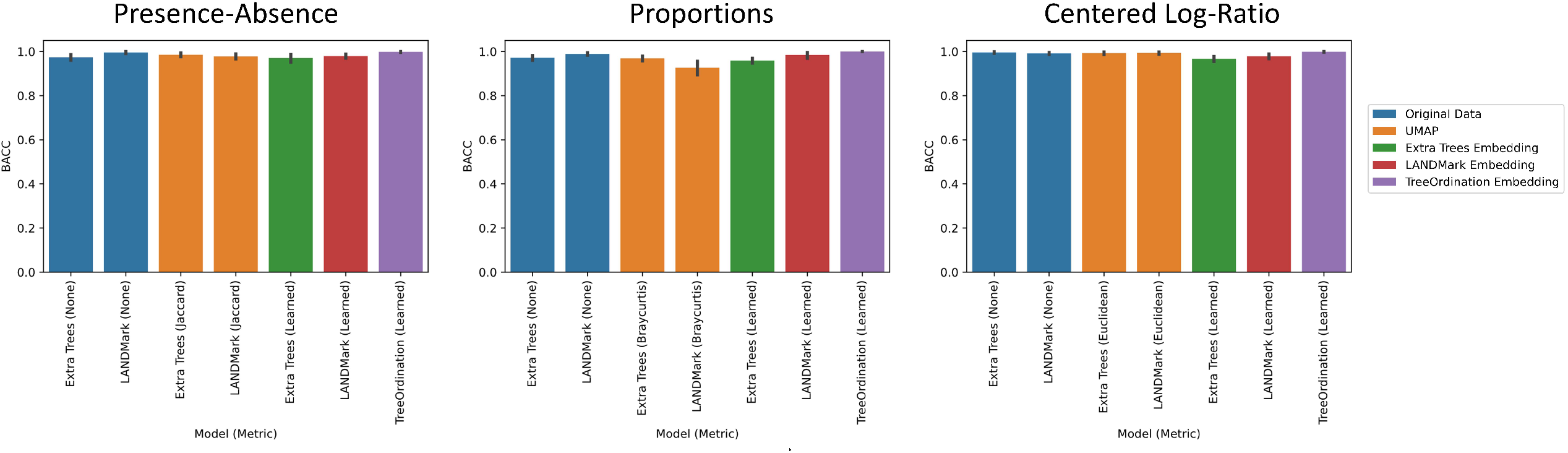
Balanced accuracy score results for each test condition in the positive control data. Higher scores indicate a more accurate classifier. The 95% confidence interval, calculated using 2000 bootstraps, is shown for each bar in black. Five-fold stratified cross-validation with five repeats was used to generate this data.

**Figure 3:**
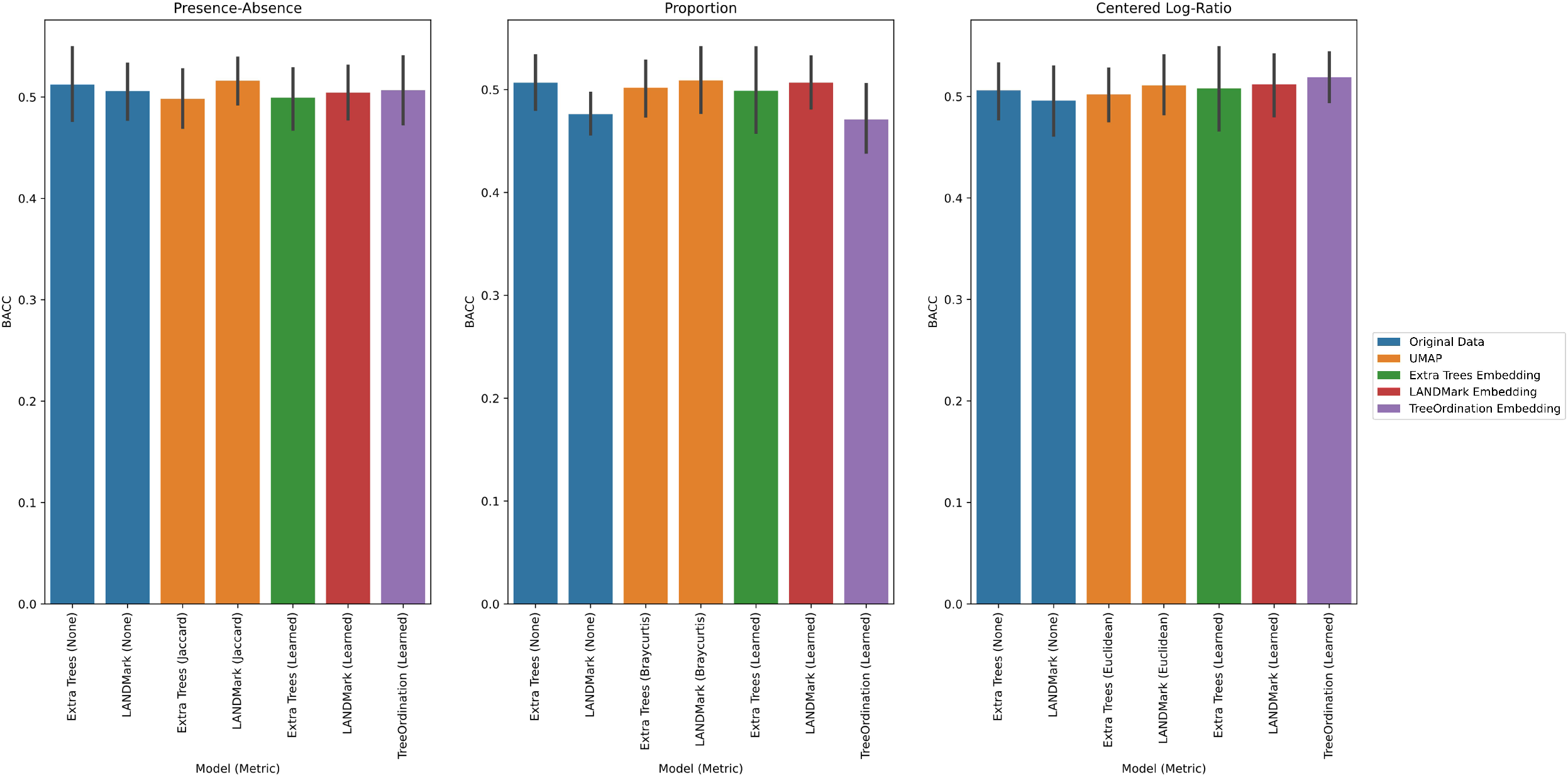
Balanced accuracy score results for each test condition in the negative control data. Higher scores indicate a more accurate classifier. The 95% confidence interval, calculated using 2000 bootstraps, is shown for each bar in black. Five-fold stratified cross-validation with five repeats was used to generate this data.

### Learning Dissimilarities Can Result in Well-Separated Groups in the Crohn’s Disease Case Study

In this study, we used the Crohn’s Disease dataset as a follow-up real-world test. In this test the CLR transformation consistently resulted in large F-statistics and significant p-values (p ≤ 0.05) (Figure 4). Except with CLR transformed data, Unsupervised Extremely Randomized Trees underperformed and often resulted in ordinations where differences between Crohn’s Disease patients and healthy controls could not be consistently detected. Even in the best case (with the CLR transform), the F-statistics associated with Unsupervised Extremely Randomized Trees ordinations was lower than those produced using the other ordination methods. We also observed that transforming data into proportions resulted in poor quality ordinations. Although reasonable ordinations were created when using the Bray-Curtis dissimilarity, the F-statistics associated with these ordinations was lower than those produced using the presence-absence and CLR transformations (Figure 4). The cross-validated generalization performance of models trained on presence-absence and CLR transformed data was the best (Figure 5). No statistically significant difference was observed in the generalization performance between models trained on CLR transformed data. However, statistically significant differences were observed between models trained on presence-absence data, proportions, and on UMAP transformed data (Supplementary Figure 3). Balanced accuracy scores calculated using TreeOrdination models were at least as good as those produced using LANDMark and Extra Trees. This result is surprising because the predictive model in TreeOrdination is trained using a high-dimensional embedding learned by at least one unsupervised LANDMark classifier.

**Figure 4:**
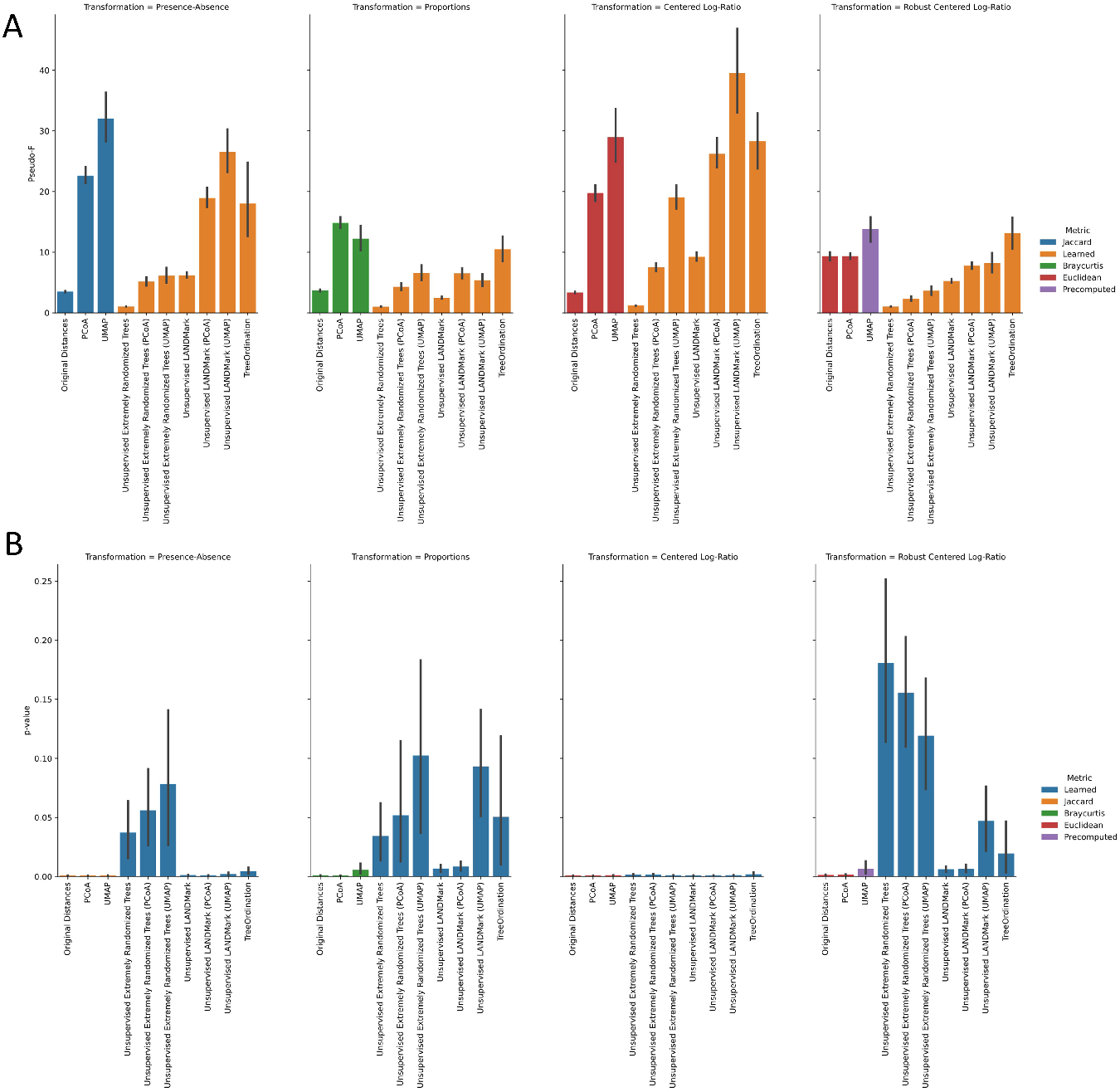
PerMANOVA results for each test condition in the Crohn’s Disease data. Higher pseudo-F statistics indicate greater separation between groups of samples (A). Panel (B) shows the distribution of p-values associated with the cross-validated distribution of pseudo-F statistics from (A). The 95% confidence interval, calculated using 2000 bootstraps, is shown for each bar in black. Five-fold stratified cross-validation with five repeats was used to generate this data.

**Figure 5:**
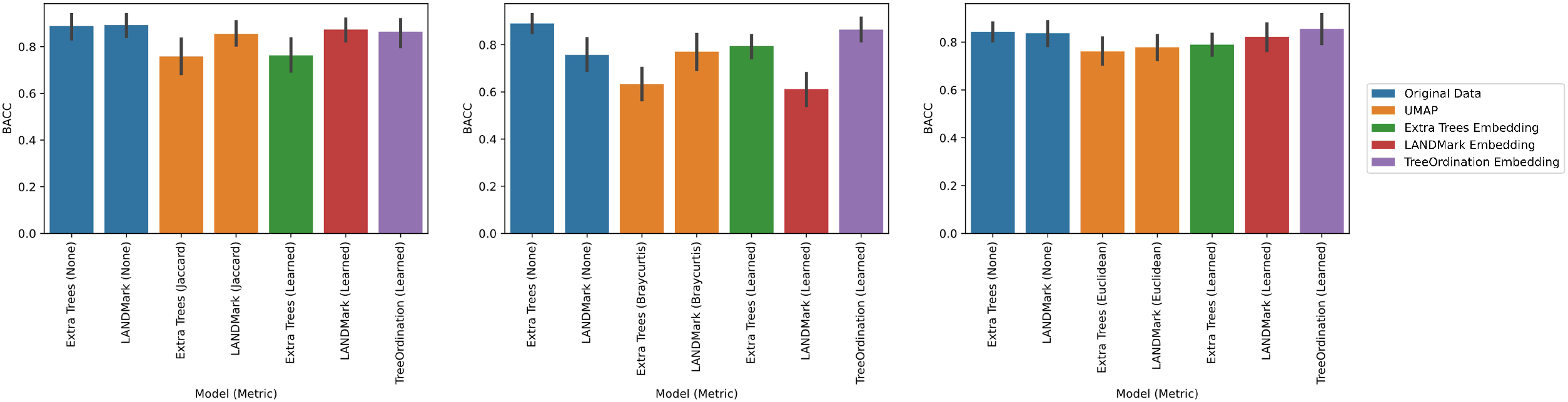
Balanced accuracy score results for each test condition in the Crohn’s Disease data. Higher scores indicate a more accurate classifier. The 95% confidence interval, calculated using 2000 bootstraps, is shown for each bar in black. Five-fold stratified cross-validation with five repeats was used to generate this data.

### LANDMark and TreeOrdination Can Be Used With Datasets Where Smaller Effect Size Exist Between Conditions

Models in this section we will adopt the naming convention for models used in Baxter et al. (2016). Specifically, classifiers trained on the combined ASV and FIT data will be called ASV Multitarget Microbiota Test (MMT) models. All models trained to distinguish between colorectal cancer and healthy controls performed better when incorporating FIT results with the microbiome data (Figure 6). All models were better than random (p ≤ 0.0001) and LANDMark and TreeOrdination ASV MMT models performed particularly well in this test and our average ROC-AUC scores for each model 0.953 and 0.953, respectively. These performed significantly better than Extra Trees and Random Forest Classifiers (p ≤ 0.0001) and the ROC-AUC score from both models is comparable with that from the original study (0.952) (28). However, using ROC-AUC scores as a measure of generalization performance can be optimistic when used with models trained on imbalanced datasets (such as this one). To address this problem, we used the balanced accuracy score since it is a better measure with imbalanced data. We found that scores produced by LANDMark and TreeOrdination ASV MMT models (mean balanced accuracy score of 0.88) are better than that of competing methods used here (Figure 6) and that this difference was statistically significant (p ≤ 0.0001). Since the TreeOrdination and LANDMark models performed best, we used them to calculate the discrimination threshold which maximizes the detection of colorectal cancer while minimizing the detection of normal tissue. These thresholds were 0.30 for LANDMark and 0.43 for TreeOrdination. When using these thresholds, both ASV MMT models discovered a substantial fraction of additional colorectal cancers (Figure 7 and Table 2) with the LANDMark model having a higher sensitivity when compared to FIT and TreeOrdination at a cost of a lower specificity (Table 2). While detecting colorectal cancer is important, it is also important to detect pre-cancerous adenomas. Therefore, we trained a model to distinguish between lesions and normal tissue. We found that statistically signficant differences exist between the generalization performance of various models and models that included FIT results performed better (Figure 8). When training each classifier using the combined ASV and FIT-score data, we observed that the balanced accuracy scores of Random models did not differ significantly from Random Forests or Extra Trees. However, both LANDMark and TreeOrdination ASV MMT models performed significantly better than Extra Trees (p ≤ 0.0001) and Random Forests (p ≤ 0.0001) classifiers trained on the same data. Both models resulted in a mean balanced accuracy scores of 0.63. Once again, we calculated the optimal discrimination threshold for distinguishing lesions (adenomas and colorectal cancers) from normal tissue. This threshold was found to be 0.69 for LANDMark and 0.66 for TreeOrdination. Compared to FIT alone, the ASV MMT models resulted in better detection of additional colorectal cancers and adenomas (Table 2). Interestingly, while both LANDMark and TreeOrdination models performed well they differed in their ability to identify different types of lesions with TreeOrdination trading sensitivity for increased specificity and the opposite occurring in LANDMark (Figures 9).

**Table 2:**
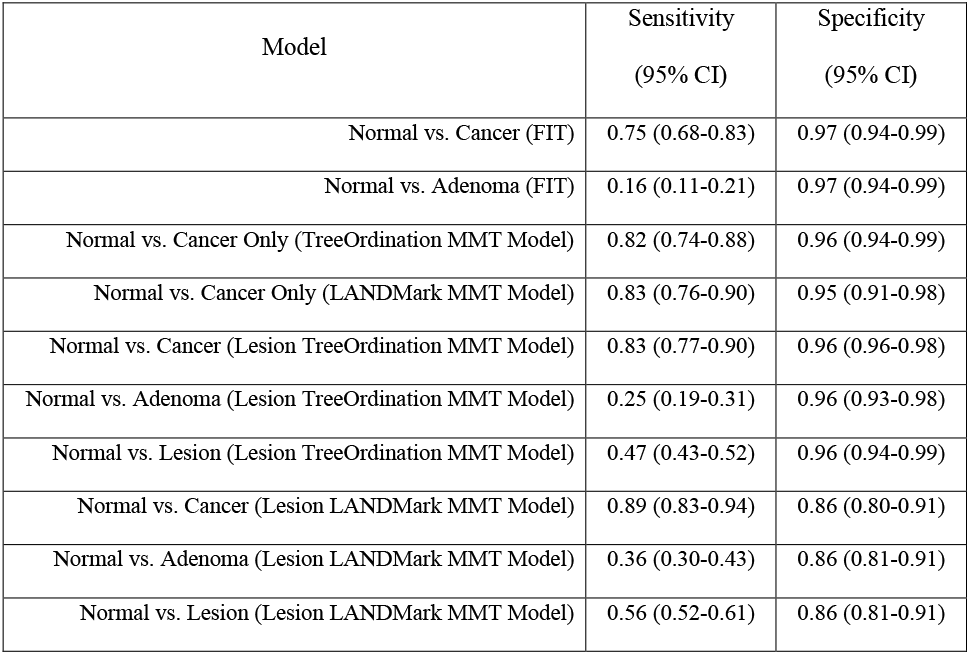
Sensitivities and specificities for LANDMark and TreeOrdination ASV MMT models, TreeOrdination ASV MMT models, and FIT. Each 95% confidence interval was calculated using 2000 stratified bootstrap replicates.

**Figure 6:**
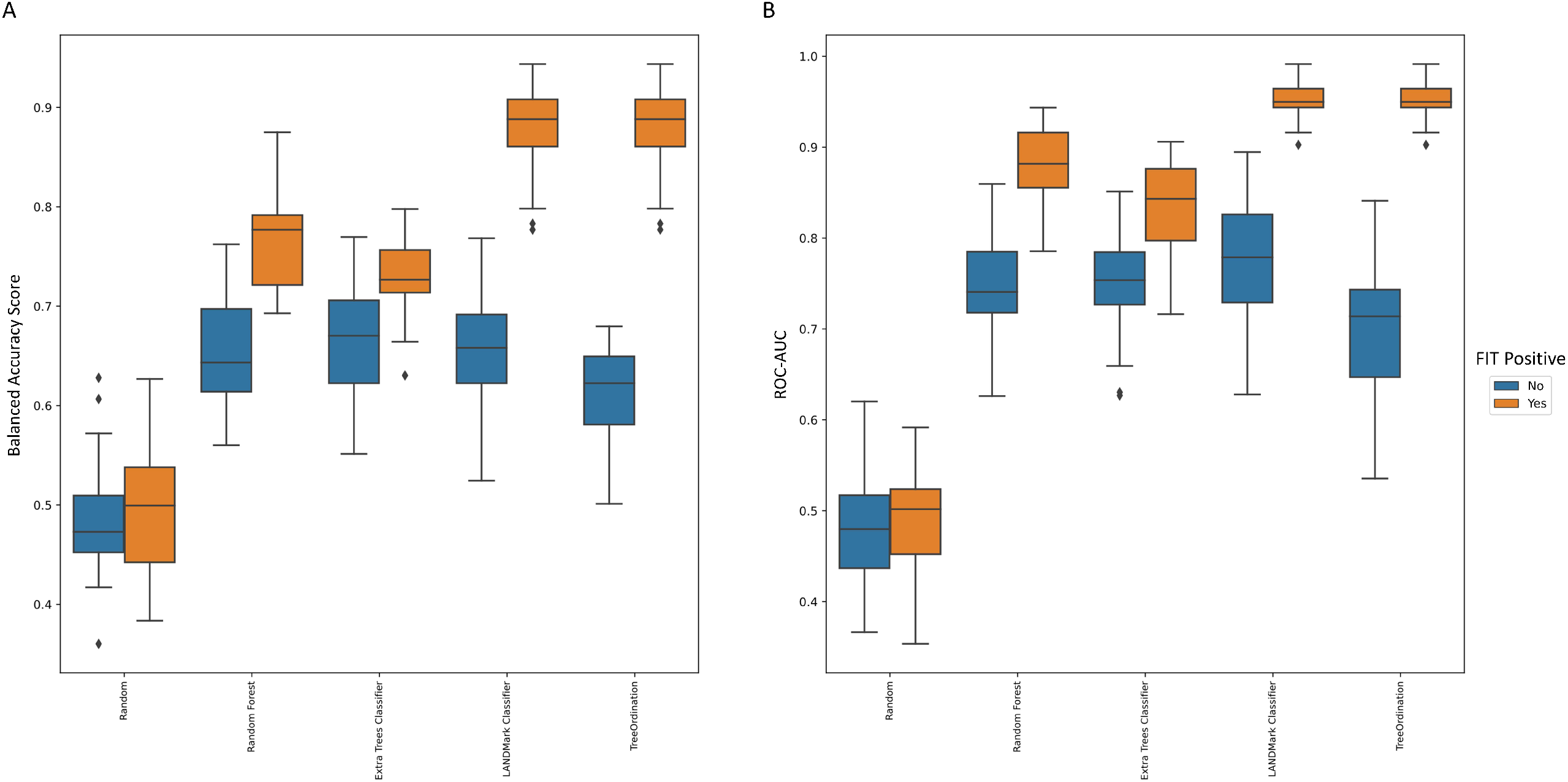
The ability of each model to identify patients with colorectal cancer. The balanced accuracy scores are reported on the left while the ROC-AUC scores are reported on the right. Each model was trained using centered log-ratio transformed data with (blue bars) and without (orange bars) the inclusion of FIT scores. These scores were generated using Fivefold Stratified Cross Validation with five repeats.

**Figure 7:**
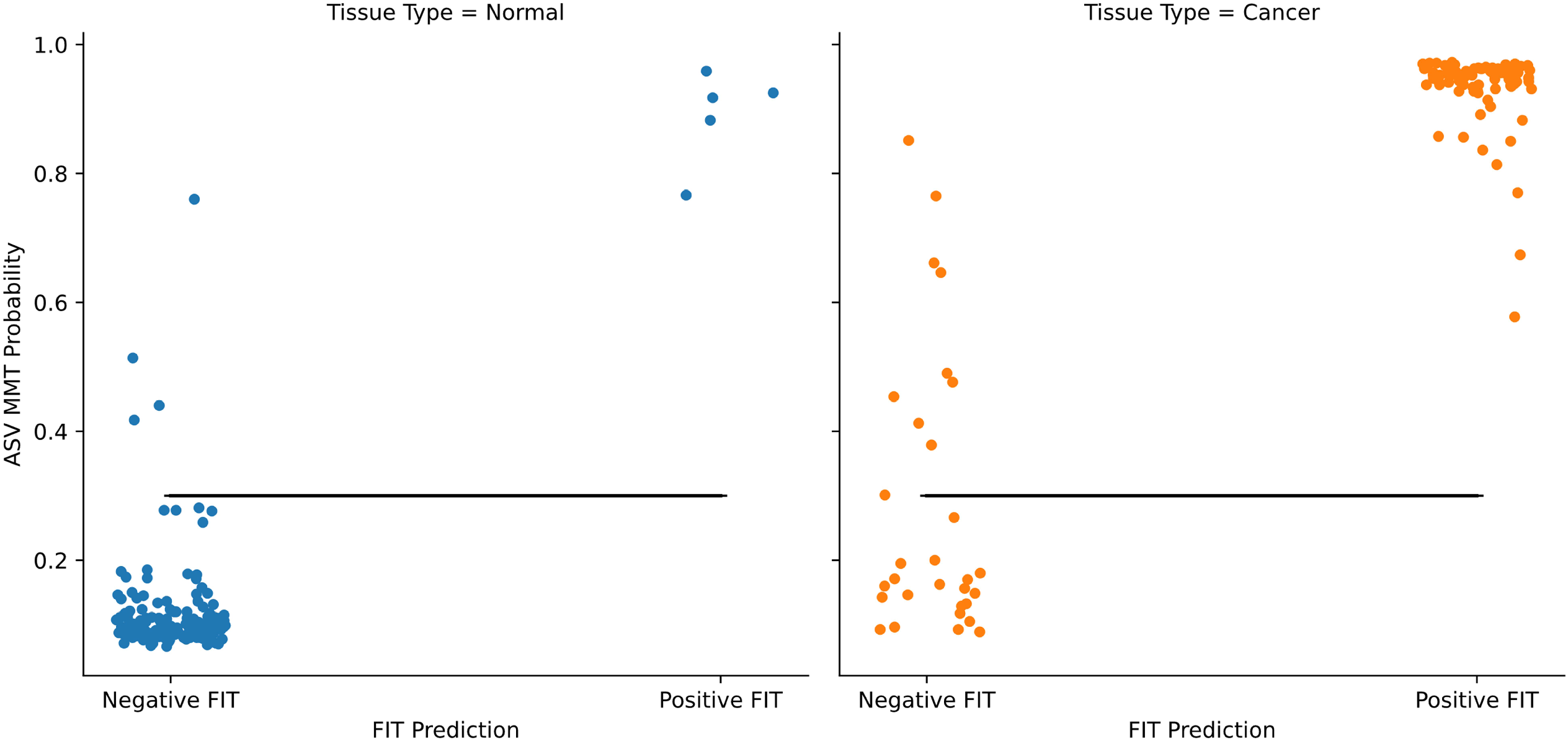
Results of the LANDMark-based ASV MMT model for distinguishing between normal and colorectal cancer. Points are colored based on the predicted labels and actual labels. The position of points along the y-axis was found by averaging the cross-validated probabilities of test samples. The black horizontal line represents the threshold maximizing the balanced accuracy score (0.30).

**Figure 8:**
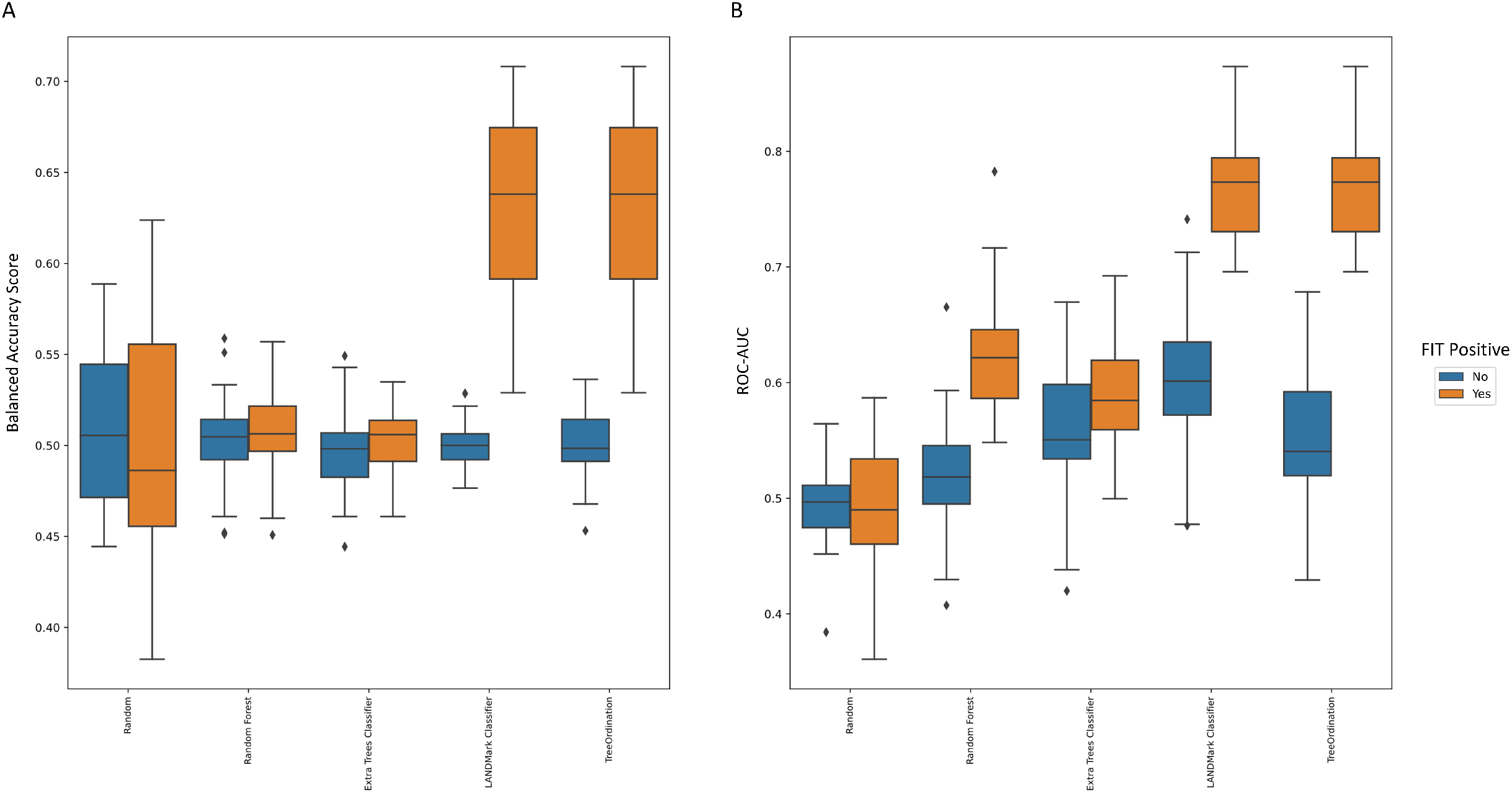
The ability of each model to identify patients with colorectal lesions (adenomas and colorectal cancers). The balanced accuracy scores are reported on the left while the ROC-AUC scores are reported on the right. Each model was trained using centered log-ratio transformed data with (blue bars) and without (orange bars) the inclusion of FIT scores. These scores were generated using five-fold stratified cross-validation with five repeats. LANDMark and TreeOrdination models trained on the combined ASV and FIT data tended to perform better than their counterparts in this task.

**Figure 9:**
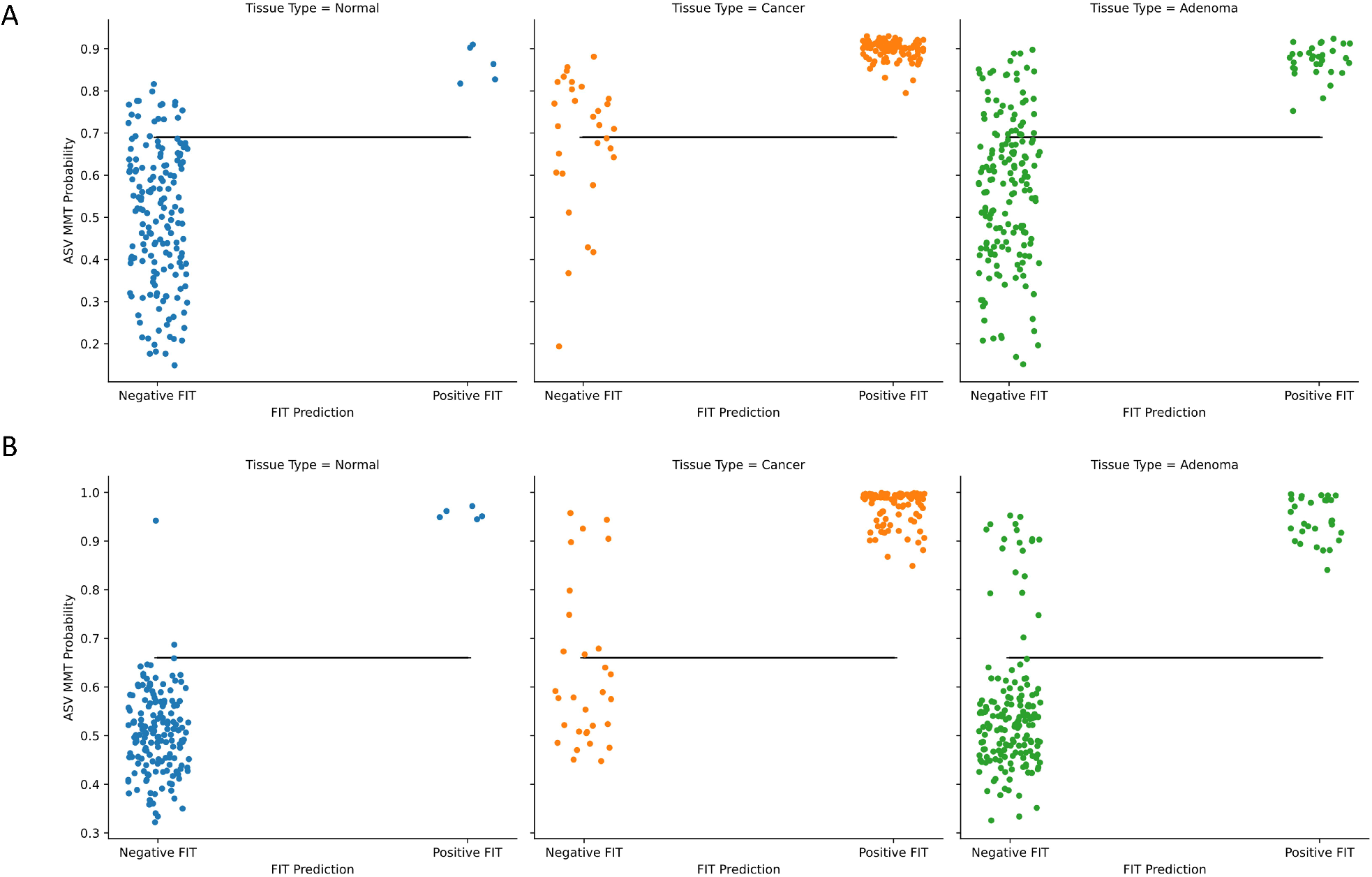
The discriminatory power of a LANDMark and TreeOrdination-based ASV MMT models. (A) The ability of LANDMark to distinguish between normal tissue, adenomas, and colorectal cancers at the given discrimination threshold (0.69). (B) The ability of TreeOrdination to distinguish between normal tissue, adenomas, and colorectal cancers at the given discrimination threshold (0.66). The position of points along the y-axis was found by averaging the cross-validated probabilities of test samples. The black line represents the threshold maximizing the balanced accuracy score (0.69).

### TreeOrdination Can be Used to Identify Features Which Have a High Impact on Model Performance

In the colorectal cancer dataset, the best discrimination threshold for a TreeOrdination ASV MMT model was found to be 0.67. While the ability of the model to identify lesions was lower than LANDMark (Table 2), this came with a marked improvement in specificity (Table 2, Figure 9). A TreeOrdination projection, using the approximate embedding function, of predicted lesions (samples over the discrimination threshold) and normal samples was created using the test set data (Figure 10A). This transformation was a reasonable projection since the mean-squared error between this approximation and the full embedding was small (1.1). Clear differences in the location of lesions and normal tissue can be seen in the plot. This suggests that there are differences between patients with colorectal lesions and those with normal tissue. This observation is supported by way of a significant PerMANOVA (pseudo-F = 375.12, p ≤ 0.001, R^2^ = 0.99). Most of the variation between samples, approximately 87%, can be explained by the first principal component, which appears to be associated with disease status. The variation along this component appears to be predominantly driven by FIT scores (Figures 10B, 11, and 12).

**Figure 10:**
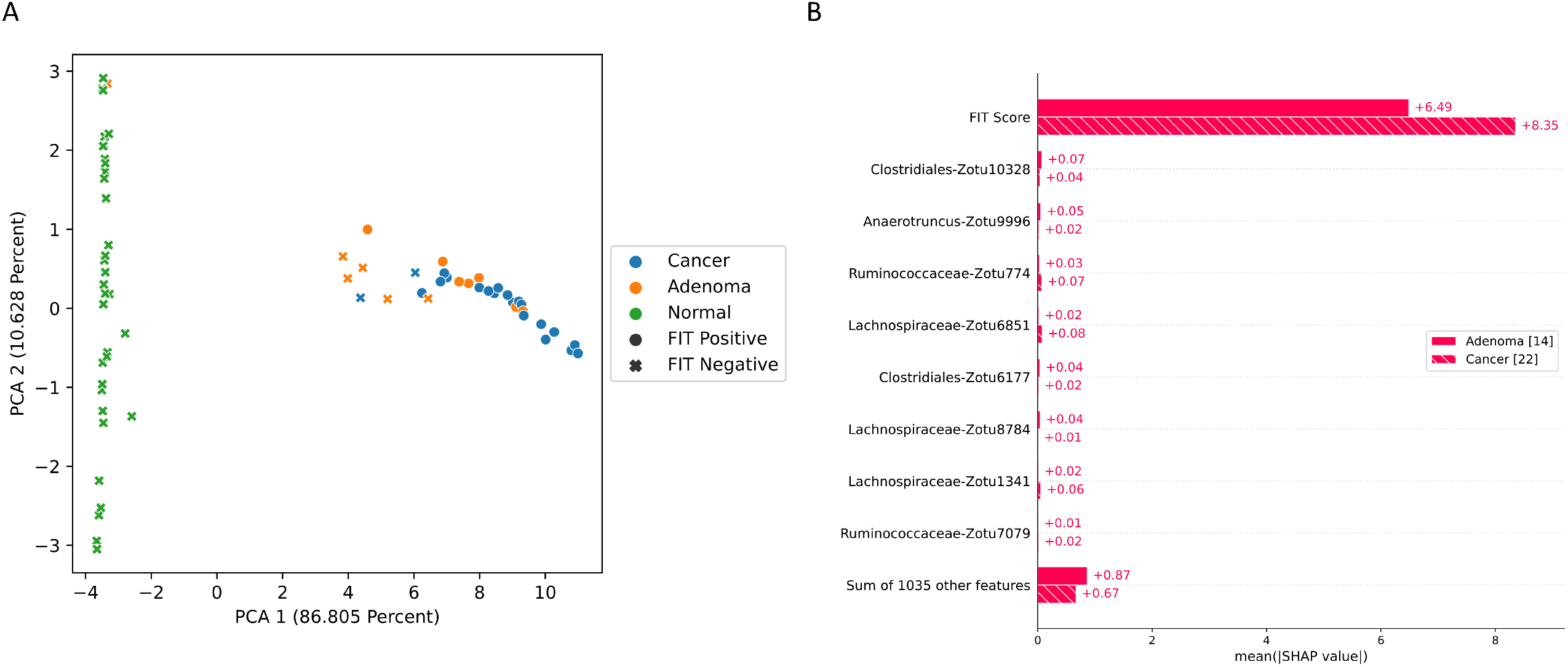
TreeOrdination projection of test samples. (A) PCA axis 1 verses axis 2 constructed using the subset of lesions (colorectal cancers and adenomas) above the discrimination threshold and all normal samples below this threshold. (B) A summary of how the Shapley values impact the placement of adenomas and carcinomas along the first principal component.

**Figure 11:**
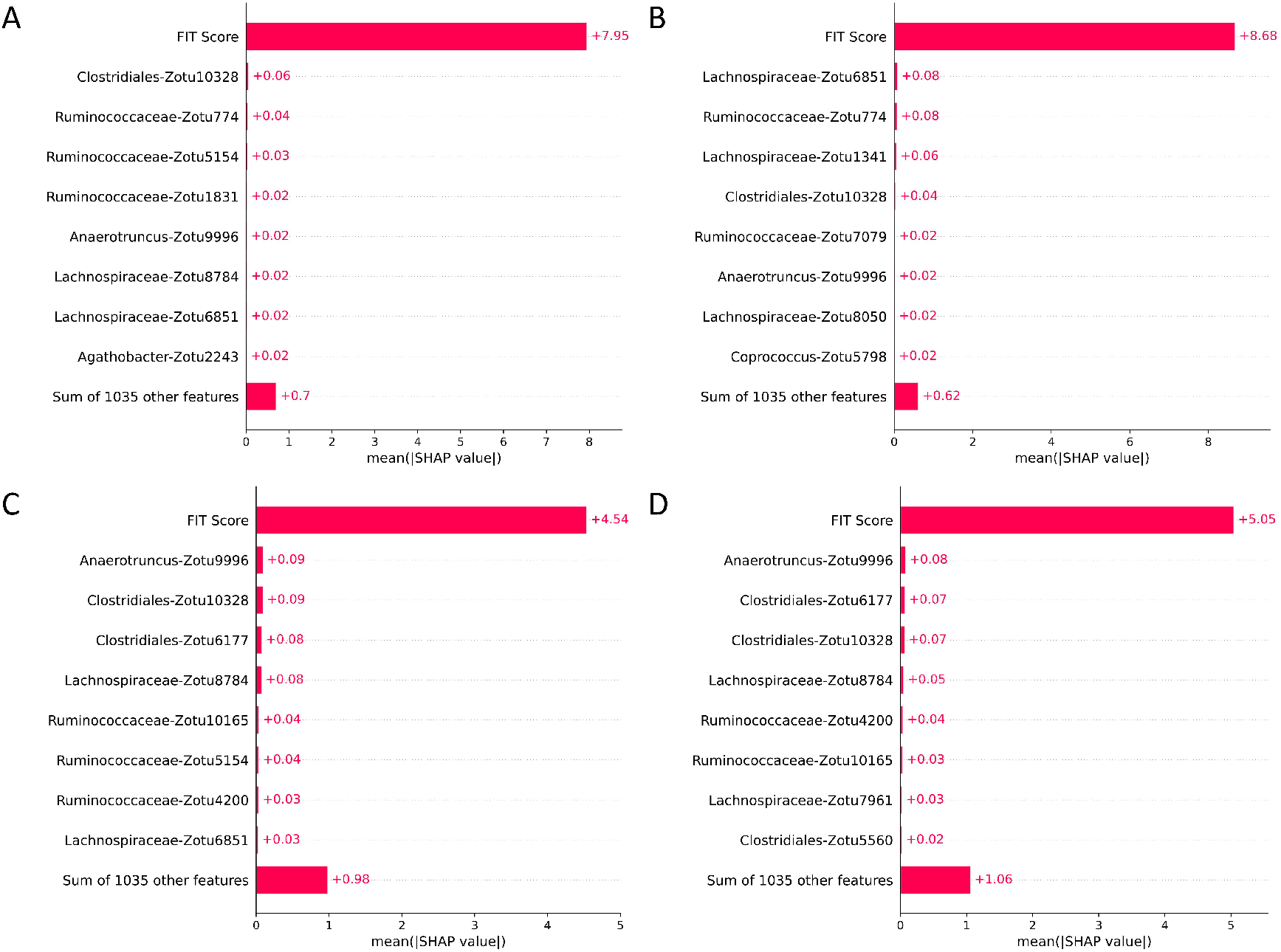
Global feature impact scores in FIT positive and FIT negative samples. The impact of each feature’s ability to discriminate the location of samples along PCA1 in the TreeOrdination projection of a FIT-positive adenomas (A), FIT-positive colorectal cancers (B), FIT-negative adenomas (C), and FIT-negative colorectal cancers (D).

**Figure 12:**
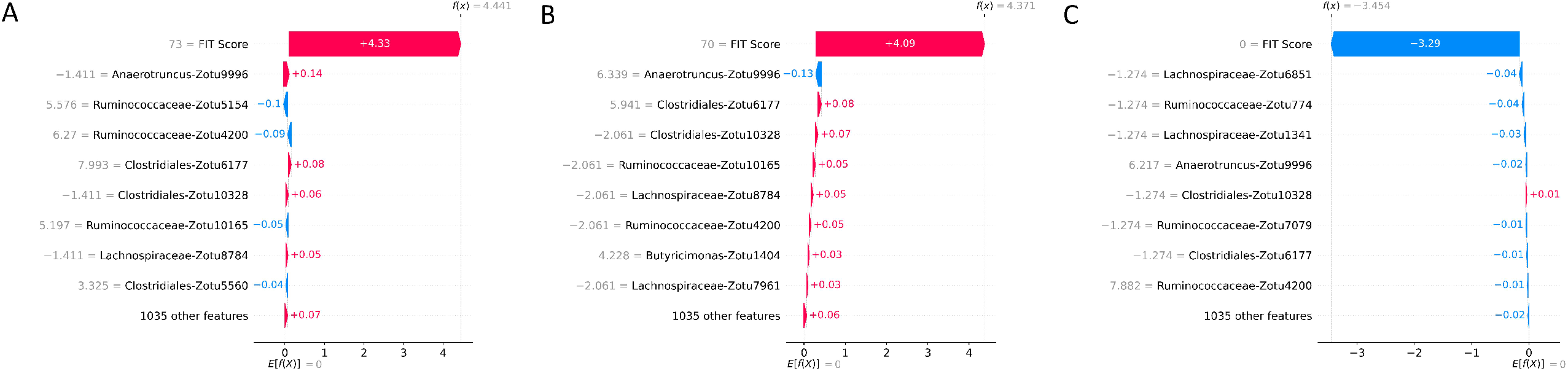
Local importance values for an individual adenoma, colorectal cancer, and normal FIT negative samples. The placement of individual samples along the first principal component (the f(x) value) can be decomposed using Shapley values. An example of how ASVs and FIT scores impact the locations of an adenoma (A), colorectal cancer (B), and a normal sample (C) are shown. Along the left in grey is the FIT score and the abundance of each ASV relative to the geometric mean of the sample. Points are shifted to the left (blue) if they are pushed towards the “normal” region of the ordination and to the right (pink) if they are pushed to the region concentrated with lesions.

Statistically significant differences were detected in the projection of adenomas and carcinoma test samples (pseudo-F = 9.51, p ≤ 0.002, R^2^ = 0.73) with variation between these groups being driven by FIT score (Figure 10B). However, differences in the microbiome also appear to play a role in determining where these samples are placed. While the impact of the microbiome is minor compared to FIT scores, ASVs assigned to *Lachnospiraceae, Clostridiales, Anaerotruncus*, and *Ruminococcaceae* are among the top features contributing to the placement of adenomas and carcenomas. Although the microbiome appears to play a more muted role, the sum of the Shapley scores for ASVs in FIT negative lesions tend to carry greater weight than their FIT positive counterparts. This could indicate that FIT scores are used to determine the region in which a sample is found while the composition of the microbiome is used to refine the location of the sample within that region. Evidence for this comes from the higher Shapley scores assigned to ASVs in FIT-negative samples (Figures 11 and 12).

The mean-squared error between the full and approximate TreeOrdination projections created using the Crohn’s Disease dataset was 0.54, indicating that the approximate embedding is reasonable. Differences between the stool microbiome of Crohn’s disease patients and healthy controls were detected using PerMANOVA (pseudo-F = 31.06, p ≤ 0.003, R^2^ = 0.99). Nearly all of the variation between samples, approximately 84%, can be explained by the first principal component. This strongly suggests that disease status is associated with the variation along this component. The ASVs which are predominantly associated with this variation belong to *Fournierella spp, Staphylococcus spp, Dialister spp*, and *Ruminococcaceae* (Figure 13).

**Figure 13:**
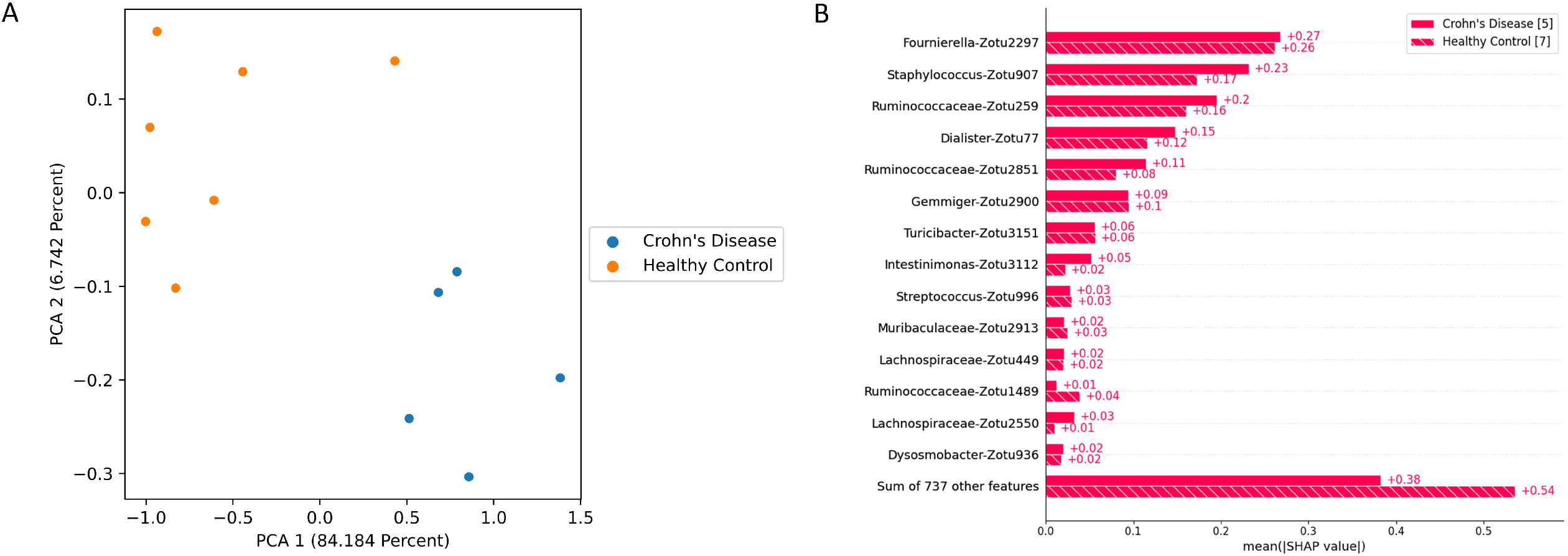
TreeOrdination projection of test samples from the Crohn’s Disease dataset. (A) PCA axis 1 verses axis 2 constructed using test samples from the Crohn’s Disease dataset. (B) A summary of how the Shapley values impact the placement of samples along the first principal component.

## Discussion

The datasets investigated here were chosen since the human gut microbiome is an important area of medical research and is becoming increasingly linked to important disease phenotypes (26,28,50,51). Since machine learning models are becoming increasingly used to identify predictive features, it is important to develop models that integrate ordination, prediction, and the detection of differentially abundant taxa so that a better understanding of the system and the composition and function of the human microbiome can be developed. The choice of transformation and dissimilarity measure is often an important first consideration when investigating microbiome data. It has long been known that the choice of dissimilarity measure can influence our measurement and interpretation of the main gradients influencing the structure of communities and taxonomic similarity between pairs of samples (52,53). For example, recent investigations have demonstrated that this choice can result in misleading results due to the sparsity inherent to the data, and differences in library size and sampling (39,54,55). To combat these problems a multitude of dissimilarity measures and ordination approaches have been developed to summarize and visualize ASV differences between sites (53). However, distance metrics and other commonly used dissimilarity measures have difficulty capturing potential dependencies between ASVs. For example, the Jaccard distance simply calculates the number of shared ASVs over the total number of unique ASVs between two communities and it fails to consider if the presence or absence of one ASV influences another (16). Furthermore, when using measures that use abundance information, differences in abundance can create situations where the sites that share the same species are more dissimilar than sites that have no species in common. Finally, as the dimensionality of the dataset grows (the number of OTUs or ASVs increases) there is an increasing probability many of these dimensions add noise to the dataset (56,57). This can occur for several reasons. For example, chimeric reads, rare sequences, inefficiencies during amplification due to primer design, and bioinformatic pipeline can contribute irrelevant information that is difficult to remove (55,58). Many contemporary dissimilarity measures are often sensitive to this noise since they have no way of reducing it when the dissimilarity is being calculated (56,57). Although the application of transformations, such as the centered log-ratio or converting to presence-absence, can mitigate some of these issues there is yet to be a consensus on which approach is best (3,39,53,59).

It is also important to address the potential weaknesses of ordination methods such as PCoA, Robust Principal Components Analysis (RPCA) (27), and UMAP. To begin, both PCoA and RPCA are unable to transform new data. This is a problem since testing and validation sets are unable to be transformed using these methods without introducing data leakage. Data leakage, in this case, can occur if projections are created using the concatenated dataset (all training, testing, and validation samples are used). When this happens, information about any withheld data is included when the PCoA or RPCA objective function is optimized. This could bias the results and potentially create machine learning models which produce overly optimistic and potentially misleading results (49). With UMAP this is not a problem since UMAP can learn an appropriate transformation using only the training data. This transformation can then be applied to test data or new data without data leakage occurring. PCoA, RPCA, and to a lesser extent UMAP are also problematic in that they require a dissimilarity measure to create an ordination. This potentially makes these methods susceptible to the previously discussed assumptions and issues that come with the use of such measures.

Our work presents results that attempt to address a gap ecology and microbial ecology – using metric learning to model and understand community structure. These approaches to measuring pairwise dissimilarity have been applied to the analysis of genomic and transcriptomic datasets (18,20,22,33). Our foray into this domain shows that learning the pairwise dissimilarities between samples can result in good projections which preserve the suspected biological differences between groups. We show that UMAP is an excellent out-of-the-box replacement for PCoA and RPCA. When using UMAP projections, we observed larger separations between groups (measured using PerMANOVA) and models trained on the projected data were accurate at identifying the class of test samples (See Figures 4-6). Further, training models using these representations results in generalization performance that is at least as good as competing methods (Figure 1 and 2). This supports a growing body of work that advocates for using UMAP to study community composition (13,60). We also attempt to address the problems associated with contemporary measures of dissimilarity (15,16,61). Specifically, we introduce an approach, TreeOrdination, which can learn the pairwise dissimilarities between samples while potentially accounting for dependencies between features since it uses a classifier, LANDMark, which makes multivariate splits at each node. In previous work, we have shown that ensembles built using such splits do a better job of modeling the underlying pattern in data (25). This is also well supported by other work in the field (15,17,24,33). By using LANDMark, TreeOrdination also minimizes the impact of noisy features through randomization (bootstrapping of training data at each node, random selection of features, and models) and regularization (most models selected for splitting are L1 or L2 regularized) (21,25). TreeOrdination also mitigates the weaknesses of PCoA and RPCA since it can transform new data after training. Furthermore, it does not require a distance metric since the dissimilarity function is learned. TreeOrdination can do this since it first creates an intermediate embedding of the training data before using this embedding as input for a UMAP projection (18,19,22,25). Once the initial embedding is learned, the training data is no longer needed since each LANDMark model has learned to create an internal representation of how to partition the space upon which the training samples are found (24,25,33,61). The location of testing samples in this space can then be extracted and transformed into a lower-dimensional space using UMAP. We show that this process works well when applied to real metagenomic datasets. In these datasets, TreeOrdination’s performance in these datasets was competitive with its peers. However, we must note that we only examined two datasets and metric learning approaches may not be suitable for all data. This is an important limitation of this study and therefore it is up to the investigator to explore potential models and use the one best suited for the data and question being asked.

A final contribution of our work is the link we identify between the original high-dimensional representation of the data to the lower-dimensional projection. Specifically, we use Shapley values to understand the impact that each feature (ASVs, OTUs, etc) has on the ordination. In ordination approaches such as PCoA, this information is exposed as loading scores along each principal component. However, these methods are limited in their ability to project data when the pattern separating groups of samples is non-linear. Furthermore, they are sensitive to outliers due to their reliance on a distance measure (56). Assuming groups can be separated using some non-linear function, the weakness of these approaches can be easily verified if low PerMANOVA F-statistics and high reconstruction losses between distances in the ordination and the original space are observed. While UMAP represents an important improvement, since it is a graph-based method that can model these non-linearities, we lose access to the loading scores and interpretability if it is used (13,35,62). To address this problem, Shapley values can be used to peer through the “black box” and quantify the impact that each feature has on the projection (37,38).

Our results support this approach since we have identified ASVs belonging to organisms associated with inflammation and colorectal cancer. This will not be an exhaustive discussion of the organisms identified since the aim of this work is to show that methods, such as TreeOrdination, are feasible. We observed that ASVs associated with *Ruminococcaceae* are important in refining the location of samples in FIT-negative adenomas and colorectal cancers and determining the location of samples in the Crohn’s Disease dataset in TreeOrdination models. The identification of this microbe is important since there is evidence to suggest that the depletion of this family is associated with colorectal cancer and gut inflammation (63,64).

Indeed, relative to the geometric mean, the depletion of *Ruminococcaceae* in the test samples pushes them to be more lesion-like (Figure 12). Our results also found commonalities with those reported in the original study. For example, our results and those from Baxter et al. (2016) show that *Lachnospiraceae* is an important predictive component and a low abundance of this group tends to push models to be more lesion-like (28,64). *Anaerotruncus* also appeared to play a role in refining the location of FIT-negative samples. The abundance of this genus relative to the geometric mean of each sample suggests that high abundance will push samples to be less lesion-like. This is an encouraging result since work has shown that an elevated abundance of *Anaerotruncus*, a butyrate producer, may exert a protective effect by inhibiting glucose transport and glycolysis (65,66). Additional support for our method comes from the results of the Crohn’s disease dataset. *Fournierella spp, Staphylococcus spp, Dialister spp*, and *Ruminococcaceae*. were identified as the organisms with the strongest impact on the location of samples in the TreeOrdination projection. We also identified ASVs assigned to *Ruminococcaceae, Gemmiger spp*., and *Lachnospiraceae* as those which impact the location of samples in the TreeOrdination projection. Encourgingly, these were also detected as important in original Forbes et al. (2018) study (26). Some taxa, such as *Staphylococcus* and *Dialister*, have known associations with Crohn’s Disease, pro-inflammatory phenotypes, and other inflammatory diseases such as Ankylosing Spondylitis (67–69). In addition, detecting ASVs assigned to *Ruminococcaceae* and *Lachnospiraceae* as impactful is important validation since these are families are known to be beneficial due to be producers of anti-inflammatory metabolites (28,70). We do acknowledge that our investigation differed from the original investigations due to our use of ASVs. However, our results are generally consistent with the wider interpretation of these studies: the composition of the microbiome tends to shift so toward species that exert a pro-inflammatory effect (26,28). Finally, these results demonstrate the power of TreeOrdination since the factors contributing to the placement of individual samples can be investigated. This allows investigators to use a single model to investigate community composition and differential abundance and predict the properties of new samples.

## Conclusions and Future Work

Our work has shown that unsupervised LANDMark (Oracle) and TreeOrdination models can learn effective dissimilarity matrices. When paired with modern dimensionality reduction approaches, such as UMAP, the global structure of the dataset is preserved. These representations can then be combined with existing matrix factorization and model introspection algorithms to create informative ordinations. Additional work should be conducted on larger datasets to understand when metric learning should be used. Furthermore, by using larger datasets the effect of individual differences will be reduced since a larger sample of the population will be used in each test set (50). This will also be a good opportunity to conduct additional comparisons between other commonly used transforms, dissimilarity measures, and read clustering approaches. Finally, with respect to TreeOrdination, additional work needs to be conducted in order to identify significantly different features. While there is still work to do, our results support the use of metric learning as a replacement for traditional dissimilarity measures in ecological research. These are advantageous since they can be used to limit the influence of outliers and uninformative features. In the datasets used here we show there is a shift towards a pro-inflammatory microbial community in the stool of patients suffering from Crohn’s disease and colorectal cancer and that our results are similar to those found elsewhere in the literature. Finally, additional work should be conducted to determine more optimal ways of calculating the dissimilarity between samples when using decision tree ensembles. This will hopefully lead to improved ordinations and interpretations.

## Acknowledgments

We would like to thank Dr. Katie McGee and Dr. Terri M. Porter for their thoughtful discussions during the development of LANDMark. We would like to thank Dr. Marko Rudar for his thoughtful review of the manuscript. Finally, we thank our reviewers and editor for their time and comments. Their work has resulted in a considerably improved manuscript.

## Supporting Information

**Supplementary File 1 – Supplementary Figure 1: High-level overview of how TreeOrdination produces a projection**. (A) Columns in a copy of the training data are randomized. The randomized data is then concatenated to the original data. (B) *N* LANDMark classifiers are trained on *N* unique randomizations of the original data. No information about the original sample labels is used to train these classifiers. (C) Using the training data, the leaf labels for each of the *N* LANDMark classifiers are extracted and concatenated together. The result is a binary matrix where each row is a sample and each column a leaf label. (D) Using this binary matrix, a UMAP/PCA transformer is trained. (E) Using the unseen or testing data, the leaf labels for each of the *N* LANDMark classifiers are extracted and concatenated together. The result is a binary matrix where each row is a sample and each column a leaf label. (F) The trained UMAP/PCA transformer from (D) is then used to transform this matrix.

**Supplementary File 2 – Supplementary Figure 2: Balanced accuracy score results for each test condition in the positive control data**. Higher scores indicate a more accurate classifier. The 95% confidence interval, calculated using 2000 bootstraps, is shown for each bar in black. Five-fold stratified cross-validation with five repeats was used to generate this data. The statistical significance of each pairwise comparison was computed using the Wilcoxon test followed by a Benjamini-Hochberg correction. p ≤ 0.0001 (****), p ≤ 0.001 (***), p ≤ 0.01 (**), p ≤ 0.05 (*), p > 0.05 (ns).

**Supplementary File 3 – Supplementary Figure 3: Balanced accuracy score results for each test condition in the Crohn’s Disease data**. Higher scores indicate a more accurate classifier. The 95% confidence interval, calculated using 2000 bootstraps, is shown for each bar in black. Five-fold stratified cross-validation with five repeats was used to generate this data. The statistical significance of each pairwise comparison was computed using the Wilcoxon test followed by a Benjamini-Hochberg correction. p ≤ 0.0001 (****), p ≤ 0.001 (***), p ≤ 0.01 (**), p ≤ 0.05 (*), p > 0.05 (ns).

**GitHub Link 1–** Raw Data and Code Used to Run the Analysis

(https://github.com/jrudar/Unsupervised-Decision-Trees)

**GitHub Link 2–** TreeOrdination Code

(https://github.com/jrudar/TreeOrdination)

**GitHub Link 3–** LANDMark Code

(https://github.com/jrudar/LANDMark)

## References

1. Silverman JD, Washburne AD S M, David LA. A phylogenetic transform enhances analysis of compositional microbiota data. eLife. 2017;6:21887.

2. Chen B, He X, Pan B, Zou X, You N. Comparison of beta diversity measures in clustering the high-dimensional microbial data. PLoS One. 2021;16(2):e0246893.

3. Weiss S, Xu ZZ, Peddada S, Amir A, Bittinger K, Gonzalez A, et al. Normalization and microbial differential abundance strategies depend upon data characteristics. Microbiome. 2017;5(27).

4. Love MI, Huber W, Anders S. Moderated estimation of fold change and dispersion for RNA-seq data with DESeq2. Genome Biology. 2014 Dec 5;15(12):550.

5. Paulson JN, Stine OC, Bravo HC, M P. Robust methods for differential abundance analysis in marker gene surveys. Nature Methods. 2013;10(12):1200–2.

6. Lin H, Peddada SD. Analysis of compositions of microbiomes with bias correction. Nature Communications. 2020 Jul 14;11(1):3514.

7. Lin H, Peddada SD. Analysis of microbial compositions: a review of normalization and differential abundance analysis. npj Biofilms and Microbiomes. 2020 Dec 2;6(1):60.

8. Segata N, Izard J, Waldron L, Gevers D, Miropolsky L, Garrett WS, et al. Metagenomic biomarker discovery and explanation. Genome Biology. 2011;12:60.

9. Elhaik E. Principal Component Analyses (PCA)-based findings in population genetic studies are highly biased and must be reevaluated. Scientific Reports. 2022 Aug 29;12(1):14683.

10. Nearing JT, Douglas GM, Hayes MG, MacDonald J, Desai DK, Allward N, et al. Microbiome differential abundance methods produce different results across 38 datasets. Nature Communications. 2022 Jan 17;13(1):342.

11. Kubinski R, Djamen-Kepaou JY, Zhanabaev T, Hernandez-Garcia A, Bauer S, Hildebrand F, et al. Benchmark of data processing methods and machine learning models for gut microbiome-based diagnosis of inflammatory bowel disease. bioRxiv [Internet]. 2021; Available from: https://www.biorxiv.org/content/early/2021/05/04/2021.05.03.442488

12. Dorrity MW, Saunders LM, Queitsch C, Fields S, Trapnell C. Dimensionality reduction by UMAP to visualize physical and genetic interactions. Nature Communications. 2020 Mar 24;11(1):1537.

13. Armstrong G, Martino C, Rahman G, Gonzalez A, Vázquez-Baeza Y, Mishne G, et al. Uniform Manifold Approximation and Projection (UMAP) Reveals Composite Patterns and Resolves Visualization Artifacts in Microbiome Data. mSystems. 0(0):e00691–21.

14. Ryo M, Rillig MC. Statistically reinforced machine learning for nonlinear patterns and variable interactions. Ecosphere. 2017;8(11):01976.

15. Cutler RD, Edwards TC, Beard KH, Cutler A, Hess KT, Gibson J, et al. Random Forests for Classification in Ecology. Ecology. 2007;88(11):2783–92.

16. Touw WG, Bayjanov JR, Overmars L, Backus L, Boekhorst J, Wels M, et al. Data mining in the Life Sciences with Random Forest: a walk in the park or lost in the jungle? Brief Bioinform. 2012;14(3):315–26.

17. Rhodes JS, Cutler A, Moon KR. Geometry- and Accuracy-Preserving Random Forest Proximities [Internet]. arXiv; 2022. Available from: https://arxiv.org/abs/2201.12682

18. Breiman L. Random Forests. Machine Learning. 2001;45(1):5–32.

19. Dalleau K, Couceiro M, Smail-Tabbone M. Unsupervised Extremely Randomized Trees. In: Phung D, Tseng VS, Webb GI, Ho B, Ganji M, Rashidi L, editors. Advances in Knowledge Discovery and Data Mining. Cham: Springer International Publishing; 2018. p. 478–89.

20. Alhusain L, Hafez AM. Cluster ensemble based on Random Forests for genetic data. BioData Mining. 2017 Dec 15;10(1):37.

21. Mentch L, Zhou S. Randomization as Regularization: A Degrees of Freedom Explanation for Random Forest Success. Journal of Machine Learning Research. 2020;21:171:1-171:36.

22. Pouyan MB, Kostka D. Random forest based similarity learning for single cell RNA sequencing data. Bioinformatics. 2018 Jul 1;34(13):i79–88.

23. Chen X, Ishwaran H. Random forests for genomic data analysis. Genomics. 2012 Jun 1;99(6):323–9.

24. Menze BH M K, Splitthoff DN K K, Hamprecht FA. On oblique random forests. In: Gunopulos D, Hofmann T, Malerba D, Vazirgiannis M, editors. Machine Learning and Knowledge Discovery in Databases. 2011. p. 453–69.

25. Rudar J, Porter TM, Wright M, Golding GB, Hajibabaei M. LANDMark: An ensemble approach to the supervised selection of biomarkers in high-throughput sequencing data. BMC Bioinformatics. 2022;23(1):110.

26. Forbes JD, Chen CY, Knox NC, Marrie RA, El-Gabalawy H, de Kievit T, et al. A comparative study of the gut microbiota in immune-mediated inflammatory diseases-does a common dysbiosis exist? Microbiome. 2018 Dec 13;6(1):221–221.

27. Martino C, Morton JT, Marotz CA, Thompson LR, Tripathi A, Knight R, et al. A Novel Sparse Compositional Technique Reveals Microbial Perturbations. mSystems. 2019 Feb;4(1).

28. Baxter NT, Ruffin MT, Rogers MAM, Schloss PD. Microbiota-based model improves the sensitivity of fecal immunochemical test for detecting colonic lesions. Genome Medicine. 2016 Apr 6;8(1):37.

29. Porter TM, Hajibabaei M. METAWORKS: A flexible, scalable bioinformatic pipeline for multi-marker biodiversity assessments. bioRxiv. 2020;

30. Rognes T, Flouri T, Nichols B, Quince C, Mahe F. VSEARCH: a versatile open source tool for metagenomics. PeerJ. 2016;4:2584.

31. Wang Q, Garrity GM, Tiedje JM, Cole JR. Naive Bayesian classifier for rapid assignment of rRNA sequences into the new bacterial taxonomy. Applied and Environmental Microbiology. 2007 Aug;73(16):5261–7.

32. Pedregosa F, Varoquaux G, Gramfort A, Michel V, Thirion B, Grisel O, et al. Scikit-learn: Machine Learning in Python. Journal of Machine Learning Research. 2011;12:2825–30.

33. Xiong C, Johnson D, Xu R, Corso JJ. Random Forests for Metric Learning with Implicit Pairwise Position Dependence. In: Proceedings of the 18th ACM SIGKDD International Conference on Knowledge Discovery and Data Mining [Internet]. New York, NY, USA: Association for Computing Machinery; 2012. p. 958–66. (KDD ‘12). Available from: https://doi.org/10.1145/2339530.2339680

34. Camargo A. PCAtest: testing the statistical significance of Principal Component Analysis in R. PeerJ. 2022;10:e12967.

35. McInnes L, Healy J, Melville J. UMAP: Uniform Manifold Approximation and Projection for Dimension Reduction. Journal of Open Source Software. 2020;3(29):861.

36. Geurts P, Ernst D, Wehenkel L. Extremely Randomized Trees. Machine Learning. 2006;63(1):3–42.

37. Lundberg SM, Lee S. A Unified Approach to Interpreting Model Predictions. In: 31st Conference on Neural Information Processing Systems (NIPS 2017 [Internet]. Long Beach; 2017. Available from: http://papers.nips.cc/paper/7062-a-unified-approach-to-interpreting-model-predictions.pdf.

38. Lundberg SM, Erion G, Chen H, DeGrave A, Prutkin JM, Nair B, et al. From local explanations to global understanding with explainable AI for trees. Nat Mach Intell. 2020;2(1):56–67.

39. Gloor GB, Macklaim JM, Pawlovsky-Glahn V, Egozcue JJ. Microbiome Datasets Are Compositional: And This Is Not Optional. Front Microbiol. 2017;8:2224.

40. Ranasinghe JA, Stein ED, Miller PE, Weisberg SB. Performance of two Southern California benthic community indices using species abundance and presence-only data: relevance to DNA barcoding. PLoS One. 2012;7(8):40875.

41. Wallen ZD. Comparison study of differential abundance testing methods using two large Parkinson disease gut microbiome datasets derived from 16S amplicon sequencing. BMC Bioinformatics. 2021 May 25;22(1):265.

42. Gloor GB, Wu JR, Pawlowsky-Glahn V, Egozcue JJ. It’s all relative: analyzing microbiome data as compositions. Ann Epidemiol. 2016 May;26(5):322–9.

43. Greenacre M, Martínez-Álvaro M, Blasco A. Compositional Data Analysis of Microbiome and Any-Omics Datasets: A Validation of the Additive Logratio Transformation. Frontiers in Microbiology [Internet]. 2021;12. Available from: https://www.frontiersin.org/article/10.3389/fmicb.2021.727398

44. team T scikit-bio development. scikit-bio: A Bioinformatics Library for Data Scientists, Students, and Developers [Internet]. 2022. Available from: http://scikit-bio.org

45. Martín-Fernández JA, Barceló-Vidal C, Pawlowsky-Glahn V. Dealing with Zeros and Missing Values in Compositional Data Sets Using Nonparametric Imputation. Mathematical Geology. 2003 Apr 1;35(3):253–78.

46. Charlier F, Weber M, Izak D, Harkin E, Magnus M, Lalli J, et al. Statannotations [Internet]. Zenodo; 2022. Available from: https://doi.org/10.5281/zenodo.7213391

47. Waskom ML. seaborn: statistical data visualization. Journal of Open Source Software. 2021;6(60):3021.

48. Hunter JD. Matplotlib: A 2D graphics environment. Computing in Science & Engineering. 2007;9(3):90–5.

49. Dong Q. Leakage Prediction in Machine Learning Models When Using Data from Sports Wearable Sensors. Comput Intell Neurosci. 2022;2022:5314671.

50. Flynn K, Ruffin MT IV, Turgeon K, Schloss PD. Spatial Variation of the Native Colon Microbiota in Healthy Adults. Cancer Prevention Research. 2018;11(7):393–402.

51. Aryal S, Alimadadi A, Manandhar I, Joe B, Cheng X. Machine Learning Strategy for Gut Microbiome-Based Diagnostic Screening of Cardiovascular Disease. Hypertension. 2020;76(5):1555–62.

52. Bacaro G, Ricotta C, Mazzoleni S. Measuring beta-diversity from taxonomic similarity. Journal of Vegetation Science. 2007;18(6):793–8.

53. Wildi O. Evaluating the Predictive Power of Ordination Methods in Ecological Context. Mathematics [Internet]. 2018;6(12). Available from: https://www.mdpi.com/2227-7390/6/12/295

54. Leite MFA, Kuramae EE. You must choose, but choose wisely: Model-based approaches for microbial community analysis. Soil Biology and Biochemistry. 2020;151:108042.

55. Hugerth LW, Andersson AF. Analysing Microbial Community Composition through Amplicon Sequencing: From Sampling to Hypothesis Testing. Frontiers in Microbiology. 2017;8:1561.

56. Zimek A, Schubert E, Kriegel HP. A survey on unsupervised outlier detection in high-dimensional numerical data. Statistical Analysis and Data Mining: The ASA Data Science Journal. 2012;5(5):363– 87.

57. Aggarwal CC, Hinneburg A, Keim DA. On the Surprising Behavior of Distance Metrics in High Dimensional Space. In: Van den Bussche J, Vianu V, editors. Database Theory — ICDT 2001. Berlin, Heidelberg: Springer Berlin Heidelberg; 2001. p. 420–34.

58. Schloss PD. Amplicon Sequence Variants Artificially Split Bacterial Genomes into Separate Clusters. mSphere. 2021 Aug 25;6(4):e0019121.

59. McKnight DT, Huerlimann R, Bower DS, Schwarzkopf L, Alford RA, Zenger KR. Methods for normalizing microbiome data: An ecological perspective. Methods in Ecology and Evolution. 2019;10(3):389–400.

60. Klimenko NS, Odintsova VE, Revel-Muroz A, Tyakht AV. The hallmarks of dietary intervention-resilient gut microbiome. npj Biofilms and Microbiomes. 2022 Oct 8;8(1):77.

61. Tsagkrasoulis D, Montana G. Random forest regression for manifold-valued responses. Pattern Recognition Letters. 2018;101:6–13.

62. Kobak D, Linderman GC. Initialization is critical for preserving global data structure in both t-SNE and UMAP. Nature Biotechnology. 2021 Feb 1;39(2):156–7.

63. He T, Cheng X, Xing C. The gut microbial diversity of colon cancer patients and the clinical significance. Bioengineered. 2021 Jan 1;12(1):7046–60.

64. Park J, Kim NE, Yoon H, Shin CM, Kim N, Lee DH, et al. Fecal Microbiota and Gut Microbe-Derived Extracellular Vesicles in Colorectal Cancer. Frontiers in Oncology [Internet]. 2021;11. Available from: https://www.frontiersin.org/articles/10.3389/fonc.2021.650026

65. Geng HW, Yin FY, Zhang ZF, Gong X, Yang Y. Butyrate Suppresses Glucose Metabolism of Colorectal Cancer Cells via GPR109a-AKT Signaling Pathway and Enhances Chemotherapy. Frontiers in Molecular Biosciences [Internet]. 2021;8. Available from: https://www.frontiersin.org/articles/10.3389/fmolb.2021.634874

66. Bianchimano P, Britton GJ, Wallach DS, Smith EM, Cox LM, Liu S, et al. Mining the microbiota to identify gut commensals modulating neuroinflammation in a mouse model of multiple sclerosis. Microbiome. 2022 Oct 17;10(1):174.

67. Huang WQ, Huang HL, Peng W, Liu YD, Zhou YL, Xu HM, et al. Altered Pattern of Immunoglobulin A-Targeted Microbiota in Inflammatory Bowel Disease After Fecal Transplantation. Front Microbiol. 2022;13:873018.

68. Lopetuso LR, Petito V, Graziani C, Schiavoni E, Paroni Sterbini F, Poscia A, et al. Gut Microbiota in Health, Diverticular Disease, Irritable Bowel Syndrome, and Inflammatory Bowel Diseases: Time for Microbial Marker of Gastrointestinal Disorders. Digestive Diseases. 2018;36(1):56–65.

69. Tito RY, Cypers H, Joossens M, Varkas G, Van Praet L, Glorieus E, et al. Brief Report: Dialister as a Microbial Marker of Disease Activity in Spondyloarthritis. Arthritis & Rheumatology. 2017;69(1):114–21.

70. Parada Venegas D, De la Fuente MK, Landskron G, González MJ, Quera R, Dijkstra G, et al. Short Chain Fatty Acids (SCFAs)-Mediated Gut Epithelial and Immune Regulation and Its Relevance for Inflammatory Bowel Diseases. Frontiers in Immunology [Internet]. 2019;10. Available from: https://www.frontiersin.org/articles/10.3389/fimmu.2019.00277

