## Supplementary figures and images for "Decision Tree Ensembles Utilizing Multivariate Splits Are Effective at Investigating Beta-Diversity in Medically Relevant 16S Amplicon Sequencing Data"

### Supplementary Figure 1

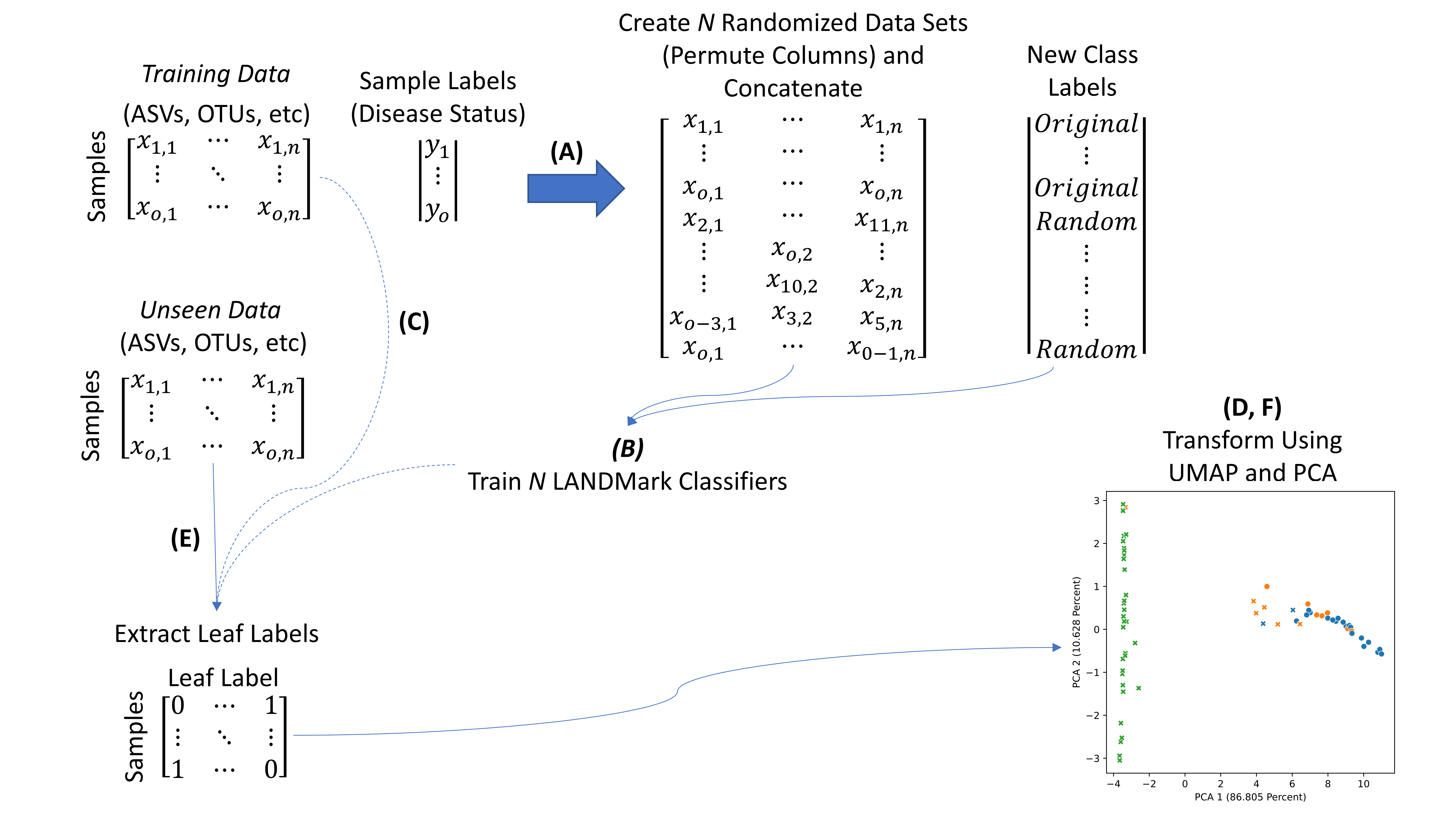

### Supplementary Figure 2

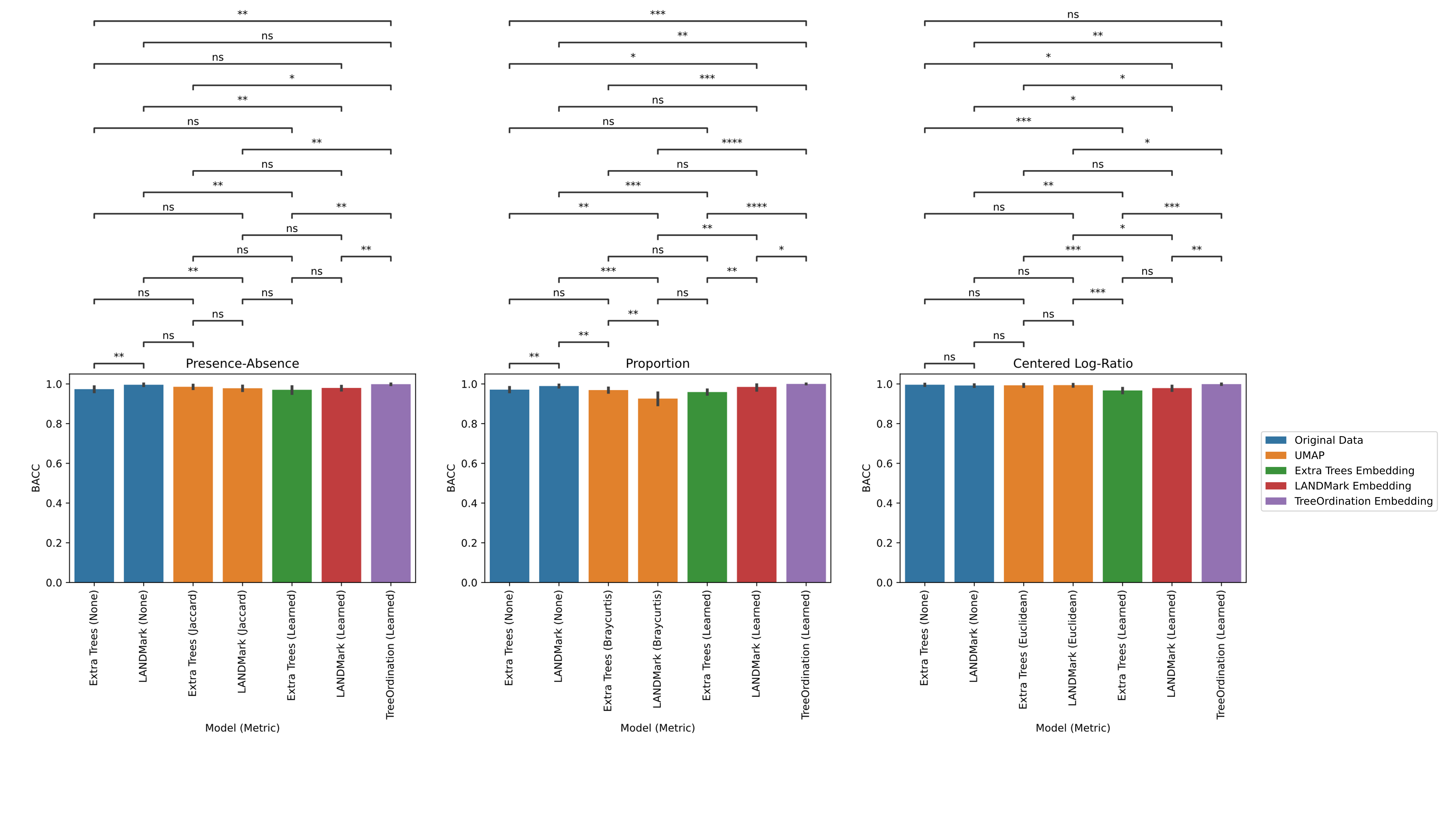

### Supplementary Figure 3

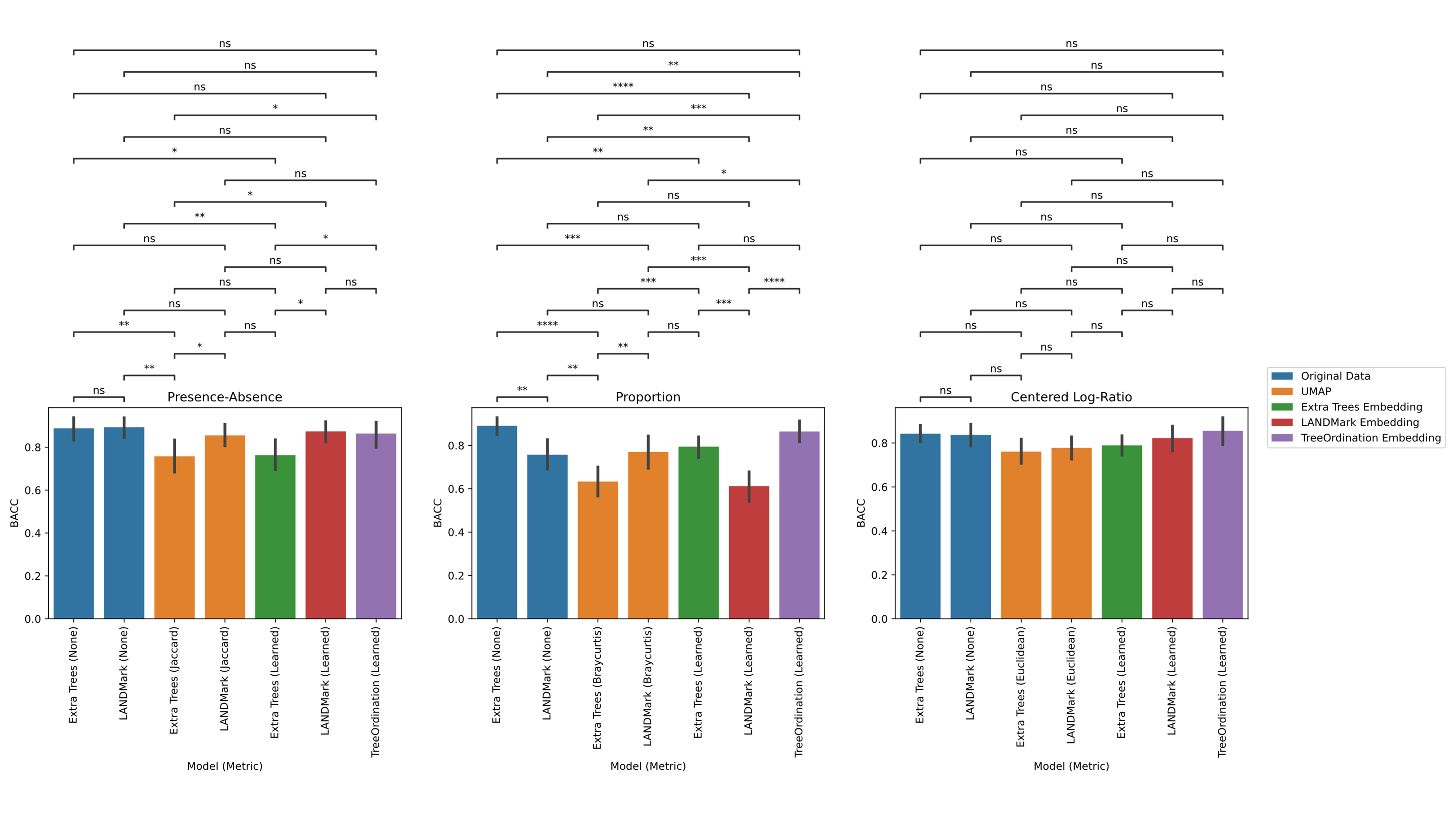
